# Three-dimensional genome re-wiring in loci with Human Accelerated Regions

**DOI:** 10.1101/2022.10.04.510859

**Authors:** Kathleen C. Keough, Sean Whalen, Fumitaka Inoue, Pawel F. Przytycki, Tyler Fair, Chengyu Deng, Marilyn Steyert, Hane Ryu, Kerstin Lindblad-Toh, Elinor Karlsson, Zoonomia Consortium, Tomasz Nowakowski, Nadav Ahituv, Alex Pollen, Katherine S. Pollard

## Abstract

Human Accelerated Regions (HARs) are conserved genomic loci that evolved at an accelerated rate in the human lineage and may underlie human-specific traits. We generated HARs and chimpanzee accelerated regions with the largest alignment of mammalian genomes to date. To facilitate exploration of accelerated evolution in other lineages, we implemented an open-source Nextflow pipeline that runs on any computing platform. Combining deep-learning with chromatin capture experiments in human and chimpanzee neural progenitor cells, we discovered a significant enrichment of HARs in topologically associating domains (TADs) containing human-specific genomic variants that change three-dimensional (3D) genome organization. Differential gene expression between humans and chimpanzees at these loci in multiple cell types suggests rewiring of regulatory interactions between HARs and neurodevelopmental genes. Thus, comparative genomics together with models of 3D genome folding revealed enhancer hijacking as an explanation for the rapid evolution of HARs.

**One-Sentence Summary:** Human-specific changes to 3D genome organization may have contributed to rapid evolution of mammalian-conserved loci in the human genome.

## Main Text

Human accelerated regions (HARs) are genomic loci that were conserved over millions of years of vertebrate evolution but evolved quickly in the human lineage, and thus are of great interest based on their potential to underlie human-specific traits (*1–8*). Many HARs are predicted to function as gene enhancers, particularly for genes implicated in neural development (*9*). Furthermore, most HARs appear to have evolved under positive selection due to having more human substitutions than expected given the local neutral rate (*10*), an indication that the sequence changes were beneficial to ancient humans. However, the mechanisms facilitating their shift in selective pressure after millions of years of constraint remains to be determined.

Structural variation is a substantial driver of genome evolution. The majority of genomic differences between humans and our closest extant relative, the chimpanzee, derive from structural variation, largely in the noncoding genome (*11*). Changes to genome organization mediated by structural variants can rewire gene regulatory networks through “enhancer hijacking”, or “enhancer adoption”, through which genes gain or lose regulatory signals, affecting spatiotemporal gene expression (*12–14*). Enhancer hijacking has been identified as a contributing factor to cancer and other human diseases (*12*, *15–17*), and previous work proposed that it may be a driver of species evolution (*7*, *18*, *19*). For example, the locus containing the cluster of Hox genes is encompassed in a single topologically associating domain (TAD) in the bilaterian ancestor, but vertebrates have two separate TADs; this difference may have driven evolutionary innovations in developmental body patterning specific to vertebrates (*18*, *20*, *21*). Recent work comparing multiple great ape genomes identified a high quality set of 17,789 human-specific structural variants (hsSVs) (*22*). We hypothesized that some HARs were hijacked due to hsSVs, changing their target gene repertoire and subjecting them to different selective pressures in humans, thus driving their human-specific accelerated evolution.

To test this hypothesis, we leverage the largest alignment of mammalian genomes to date, Zoonomia (*23*). We first identify an updated set of HARs (zooHARs) and chimpanzee accelerated regions (zooCHARs), and develop an open-source Nextflow pipeline for reproducible and streamlined identification of accelerated regions (ARs) in any lineage using large multiple sequence alignments. We find that TADs containing hsSVs are enriched for zooHARs. Using Akita, a deep learning model of three-dimensional (3D) genome folding, we predict that multiple hsSVs change the chromatin interactions of zooHARs and zooCHARs. We then validate these predictions by generating high-resolution chromatin capture (Hi-C) data from human and chimpanzee induced pluripotent stem cell derived neural progenitor cells (NPCs) at matched developmental time points and show that differentially expressed genes from NPCs (*24*) and cerebral organoids (*25*) are enriched in TADs containing zooHARs and hsSVs (Chi-squared p-value < 0.05). By integrating a machine learning model of enhancer activity, a network-based cell type labeling method, and a massively parallel reporter assay (MPRA) performed on primary cells from the human mid-gestation telencephalon, we characterize the regulatory activity of zooHARs and zooCHARs in specific neuronal cell types. Taken together, these results implicate enhancer hijacking as a genetic mechanism to explain the lineage-specific accelerated evolution of many HARs, potentially underlying human-specific neurodevelopmental phenotypes.

## Human accelerated regions are enriched in 3D topological associating domains with human-specific structural variants

The identification of species-specific accelerated regions in alignments containing many species with large genomes requires significant computational resources. Pipeline management software enables analyses like these to be made portable to different parallel computing environments (*26*). Therefore, we compiled previously developed methods for detecting accelerated regions (*1*, *27–29*) into a new Nextflow pipeline and optimized modeling parameters in the Phylogenetic Analysis with Space/Time models (PHAST) software package for large multiple sequence alignments, creating a scalable software tool for identification of lineage-specific accelerated elements in any species on any computing platform (Fig. 1A, Supplemental Text).

**Fig. 1.**
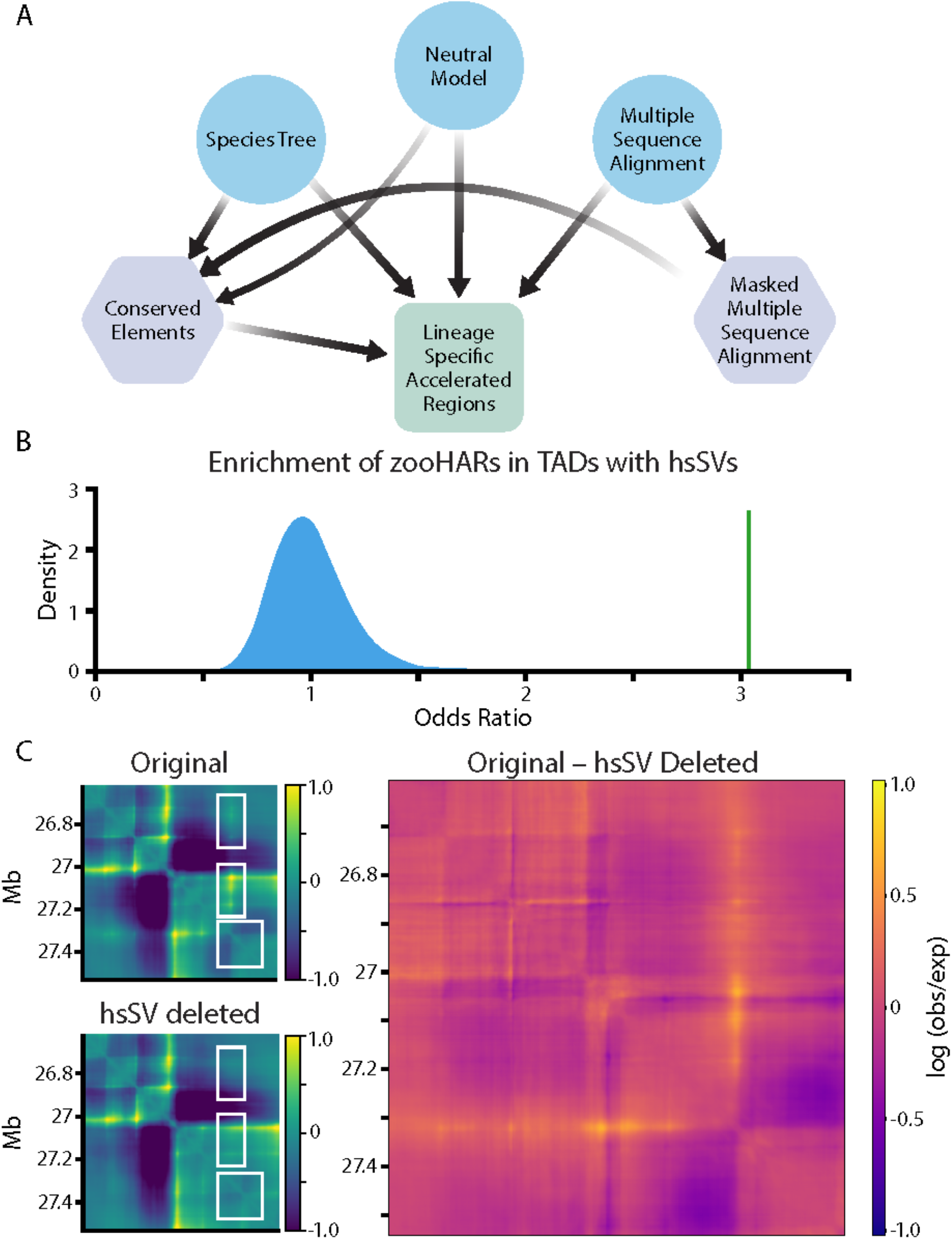
Human-specific structural variants are enriched in zooHAR chromatin domains and predicted to change the 3D genome. (**A**) Pipeline to identify lineage-specific accelerated regions. Blue circles indicate initial input data, purple hexagons are intermediate results, and the green square is the final output. (**B**) Odds ratio of chromatin contact domains in GM12878 cells (*34*) containing hsSVs and zooHARs (green line) compared to a null distribution (shaded blue region) of odds ratios for chromatin contact domains containing conserved (phastCons) elements and hsSVs from 1000 random draws of phastCons equaling the number of zooHARs. (**C**) Akita prediction of a locus (hg38.chr4:26614489-27531993, hsSV: human-specific insertion *O_000012F_1_28503465_quiver_pilon_11099913_11099913* from (*22*)) with a human-specific insertion (“Original”), with the human-specific insertion deleted *in silico* (“hsSV deleted”) and a subtraction matrix (“Original - hsSV deleted”) comparing the chromatin contact matrices with and without the human-specific insertion. White boxes indicate regions that change in the “Original” compared to the “hsSV deleted” sequences. Log(observed/expected) contact values are shown in the heatmaps.

We then leveraged the Zoonomia alignment of 241 mammal genomes to identify 312 zooHARs and 141 zooCHARs (Table S1, Table S2). These ARs demonstrate similar features to previous sets of HARs, including being mainly noncoding, having signatures of positive selection (82% of zooHARs and 86% of zooCHARs), and being located near genes involved in developmental and neurological processes (Fig. S1-3) (*6*, *9*, *10*). Approximately one-third of zooHARs and zooCHARs are transcribed in the developing human neocortex (Fig. S1E-F). The median distance between zooHARs and zooCHARs is significantly less than expected (1.05Mb, bootstrap p-value=0.02, both in hg38), as observed in previous sets of primate accelerated regions (*30*). Genes near both zooHARs and zooCHARs are significantly enriched for roles in transcriptional regulation (hypergeometric tests (*31*, *32*); Fig. S2, 3). As human and chimpanzee ARs demonstrate similar characteristics, the smaller number of zooCHARs is likely attributable to the lower quality of the chimpanzee reference genome and the strict filtering we performed, though the annotations of genes nearby zooHARs suggest connections to a broader diversity of developmental processes compared to zooCHARs. Together these analyses demonstrate that zooHARs identified from an alignment of 241 mammals demonstrate features consistent with previous studies proposing gene regulatory functionality, particularly in neurodevelopment.

Genomic loci near duplicated genes have been shown to evolve rapidly, suggesting synergy between structural variation and sequence-based genome evolution (*33*). To explore this, we sought to determine whether zooHARs and hsSVs tended to co-locate in the context of the 3D genome. Using a high-quality set of TADs from lymphoblastoid cells (*34*), we found that zooHARs are strongly enriched in TADs with hsSVs relative to the set of conserved (phastCons) elements from which zooHARs are identified (odds ratio = 3.0, bootstrap p-value < 0.001, Fig. 1B). This enrichment is robust to repeating the analysis with TADs from other cell types, including primary mid-gestation telencephalon, and a different TAD-calling method, but it is not observed with random genomic windows (Fig. S4). To determine whether the enrichment is simply driven by localization of hsSVs near zooHARs in the 1D genome sequence, we replaced the TADs with random size-matched windows and found that zooHARs were not significantly enriched in this context relative to phastCons elements (fig. S4D-E). Thus, we conclude that zooHARs are specifically enriched in TADs with hsSVs, suggesting a role for 3D genome organization and structural variation in the accelerated evolution of HARs.

## Human-specific structural variants are predicted to have changed the 3D chromatin environment of zooHARs

Structural variants are the main contributor to genome-wide genetic divergence between the human and chimpanzee genomes (*11*), and they have the potential to generate large changes in 3D genome organization through disruption of insulating boundaries or other structural motifs (*35*). Based on our observation that zooHARs are enriched in TADs with hsSVs, we sought to determine whether hsSVs may have generated changes in the 3D genome near zooHARs. Using Akita, a neural network-based machine learning model trained on six cell types to predict 3D genome contact matrices from DNA sequence (*36*), we assessed the impact of hsSVs (Table S3). For each variant, we predicted the chromatin contact matrices for the DNA sequence with and without the variant and computed the mean squared distance between the two matrices. Many hsSVs are predicted to change 3D genome organization near zooHARs and zooCHARs; 30% of zooHARs and 27% of zooCHARs occur within 500 kb of a hsSV with a disruption score in the top decile of all disruption scores for hsSVs. These results suggest that human-specific 3D genome structures are encoded in DNA sequence and modified through hsSVs.

## High-resolution Hi-C data from human and chimpanzee validates 3D genome reorganization near zooHARs and zooCHARs

In order to validate the predicted changes to 3D genome organization mediated by hsSVs near zooHARs, we generated Hi-C data from NPCs differentiated from two human and two chimpanzee induced pluripotent stem cell lines, together generating over 3.4 billion uniquely mapped chromatin contacts (Table S4)(*37*). All lines were from male individuals, and two replicates were generated per sample. Stratum-adjusted correlation coefficients (*38*) demonstrated high concordance of data between replicates and individuals from the same species (Fig. S5), so we merged data from replicates and samples from the same species for downstream analyses. The *cis/trans* interaction ratio and distance-dependent interaction frequency decay indicate that the data is high quality (Table S4, Fig. S6).

Conservation of 3D genome structures, such as A and B compartments and TAD boundaries, has been demonstrated in various species, however our understanding of the extent of this conservation is still developing (*34*, *39–44*). We found 10% of TAD boundaries to be species-specific (Table S5), slightly less than the 14% identified in a recent study comparing human and macaque chromatin organization (*42*), likely due to chimpanzees being more closely related to humans than are macaques. The majority of chromatin loops, also termed ‘dots’ or ‘peaks’ (*45*), are conserved or partially conserved (Table S5, Fig. S7) (*46*, *47*). These results support the idea of conservation of large-scale chromatin structures between human and chimpanzee, though differences are detectable in specific loci.

We next confirmed the enrichment of zooHARs in TADs containing hsSVs in our Hi-C data from human NPCs (Fig. S4C, Table S5). This enrichment was also observed between zooCHARs and chimpanzee-specific structural variants (*22*) in TADs from the chimpanzee data (odds ratio=4.8, bootstrap p-value=0.04), indicating that co-location of lineage-specific structural variants and ARs is not a human-specific phenomenon. As SVs and Hi-C data are generated for more species, it will be possible to use the tools from this study to quantify this striking association across Eukaryotes. Finally, we used our NPC Hi-C data to associate zooHARs and zooCHARs with genes and found significant enrichment for transcriptional regulators of developmental processes, confirming and extending our GO results based on nearby genes.

## Hijacked zooHARs and zooCHARs are associated with differentially expressed genes

We next used gene expression data from NPCs (*24*) and cerebral organoids (*25*) derived from human and chimpanzee induced pluripotent stem cells to test if zooHARs with altered chromatin interactions are associated with altered gene regulation. We observed that differentially expressed genes in both datasets are enriched in TADs containing zooHARs and hsSVs (chi-squared p-values < 0.05). In contrast, genes differentially expressed between human and chimpanzee adult brain tissue (*48*), induced pluripotent stem cells (iPSC), iPSC-derived cardiomyocytes, and heart tissue (*49*) are not enriched in TADs containing zooHARs and hsSVs, suggesting that the effects of enhancer hijacking may be developmental stage and cell type specific.

The loci encompassing zooHAR.126 and zooHAR.15 are two clear examples of how hsSVs can alter 3D regulatory interactions between HAR enhancers and neurodevelopmental genes. Each locus has a strong Akita prediction of altered genome folding in the presence of an hsSV, which is highly similar to the differences observed in NPC Hi-C data (Fig. 2A, B) (*36*). The average disruption peaks at specific genomic elements within the 1Mb region (Fig. 2C, D), including at species-specific loops and the promoters of genes differentially expressed between humans and chimpanzees (Fig. 2E, F). For example, the Tourette’s syndrome gene *NECTIN3 (50)* is in the same TAD with an hsSV and zooHAR.126, and it is downregulated in human versus chimpanzee NPCs (*24*). Similarly, the developmental gene *MAF*, implicated in Ayme-Gripp syndrome, is differentially expressed between human and chimpanzee in inhibitory neurons, NPCs, iPSCs, iPSC-derived cardiomyocyte progenitors (*24*, *25*, *49*), and it is in a TAD encompassing a hsSV and zooHAR.15, which overlaps previously identified 2xHAR.21 (*51*). In order to determine with higher confidence that the observed changes in 3D structure at these loci were human-derived, we assessed the orthologous loci in previously published rhesus macaque fetal brain cortex plate (*42*). For both loci, the human-specific changes to 3D genome organization described here were not observed in rhesus macaque data, suggesting that they are human-derived as a result of the hsSVs, as predicted by Akita (Fig. S8) (*36*). Together, these results establish that the 3D genome changes in these loci are human-specific, associated with gene expression changes and likely caused by the hsSVs.

**Fig. 2.**
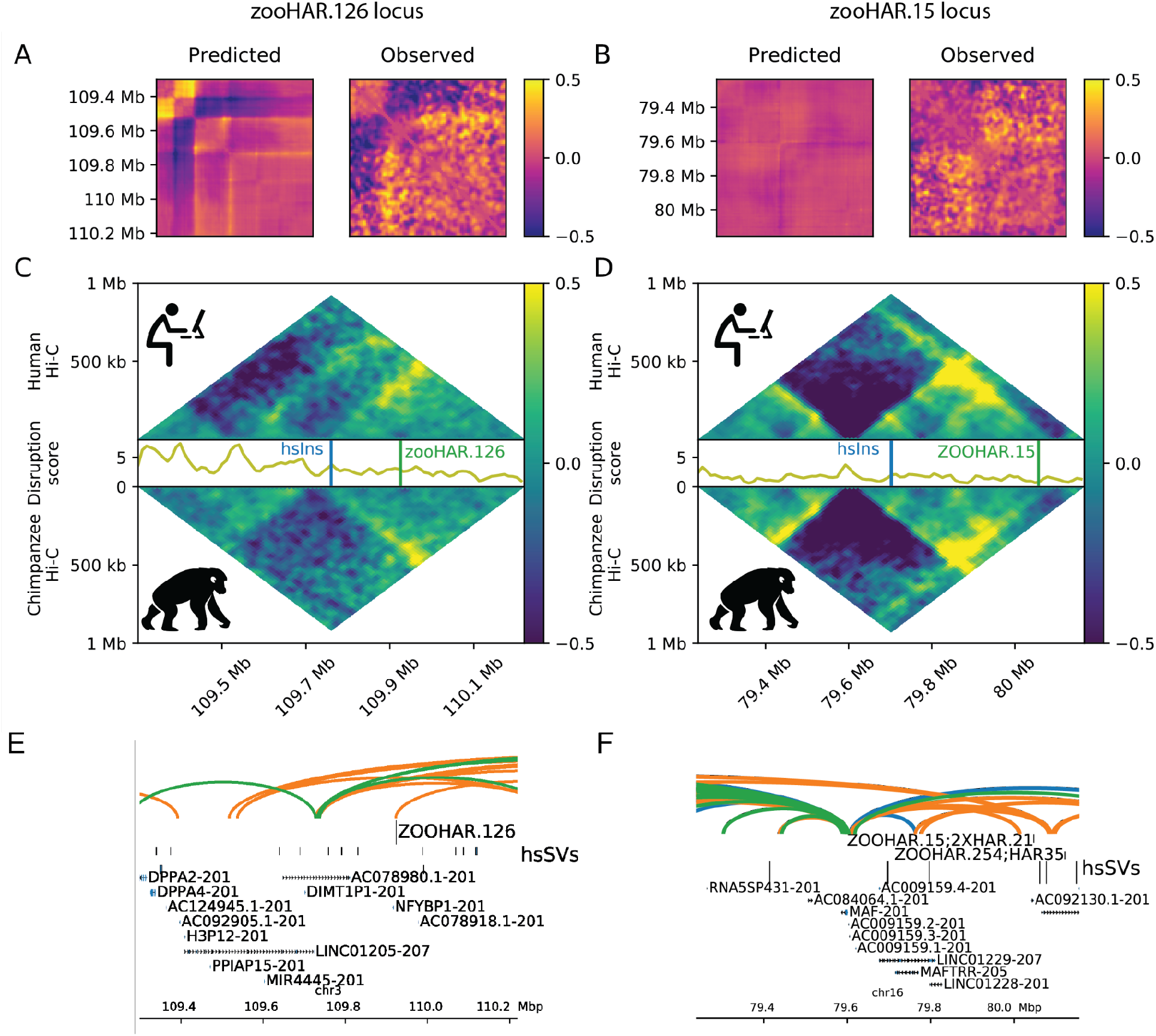
Human-specific structural variants change the 3D genome around zooHARs and zooCHARs. White boxes highlight differences between the species. Log(observed/expected) values are shown in the heatmaps. (**A, B**): Subtraction matrices for the *in silico* predicted change due to the human-specific insertion (left) and observed chromatin contact maps in human compared to chimpanzee NPC Hi-C (right) for the loci containing zooHAR.126 (hg38.chr4:26614489-27531993; hsSV: *chr4_27070203_DEL_chimpanzee_000012F_1_28503465_quiver_pilon_11099913_11099913* from (*22*))) and zooHAR.15 (hg38.chr16:79237694-80155198; hsSV: *chr16_79695894_DEL_chimpanzee_000093F_1_10181781_quiver_pilon_1690619_1690619* from (*22*)), respectively. (**C, D**): Human (top) and chimpanzee (bottom) log(observed/expected) Hi-C contact frequencies in each locus, with the disruption score (10 kilobase resolution) in between. (**E, F**): zooHAR locations denoted by vertical lines adjacent to their names. Conserved (blue), chimpanzee-specific (green), and human-specific (orange) loops (5 kilobase resolution, loops called with Mustache (*46*))

## Many zooHARs are neurodevelopmental enhancers with cell type-specific activity

In order to define the cell types and tissues that may be impacted by hijacked HARs, we expanded on previous work demonstrating enhancer-associated epigenomic signatures of HARs in specific cell types and tissues and predicting enhancer activity (*52*) by including recently generated data from 61 ATAC-seq, 40 DNase-seq and 204 ChIP-seq datasets in 44 cell types including multiple brain regions from specific developmental timepoints (*53–60*). Even against a stringent background set of phastCons elements, which themselves tend to be enriched for gene regulatory marks related to development (*9*), zooHARs are enriched for markers indicative of neurodevelopmental regulatory activity including ATAC-seq peaks and promoter capture Hi-C interactions in multiple neuronal cell types (bootstrap p < 0.05; Fig. S9). For example, zooHAR.126 overlaps numerous regulatory epigenomic marks and footprints for seven transcription factors (Fig. 3A). Over all zooHAR footprints, enriched transcription factors included inhibitory neuron specifier DLX1 (*61*), master brain regulator and telencephalon marker FOXG1, and cortical and striatal projection neuron marker MEIS2 (*62*, *63*) (Fig. 3B, Table S6). Using these datasets as features, we trained a new machine learning model on *in vivo* validated VISTA enhancers (*64*) and used it to predict that 197/312 zooHARs (63.1%) function as neurodevelopmental enhancers based on their epigenetic profiles. This increases the proportion of HARs with predicted regulatory activity in the brain relative to previous work (Table S1) (*9*, *54*).

**Fig. 3.**
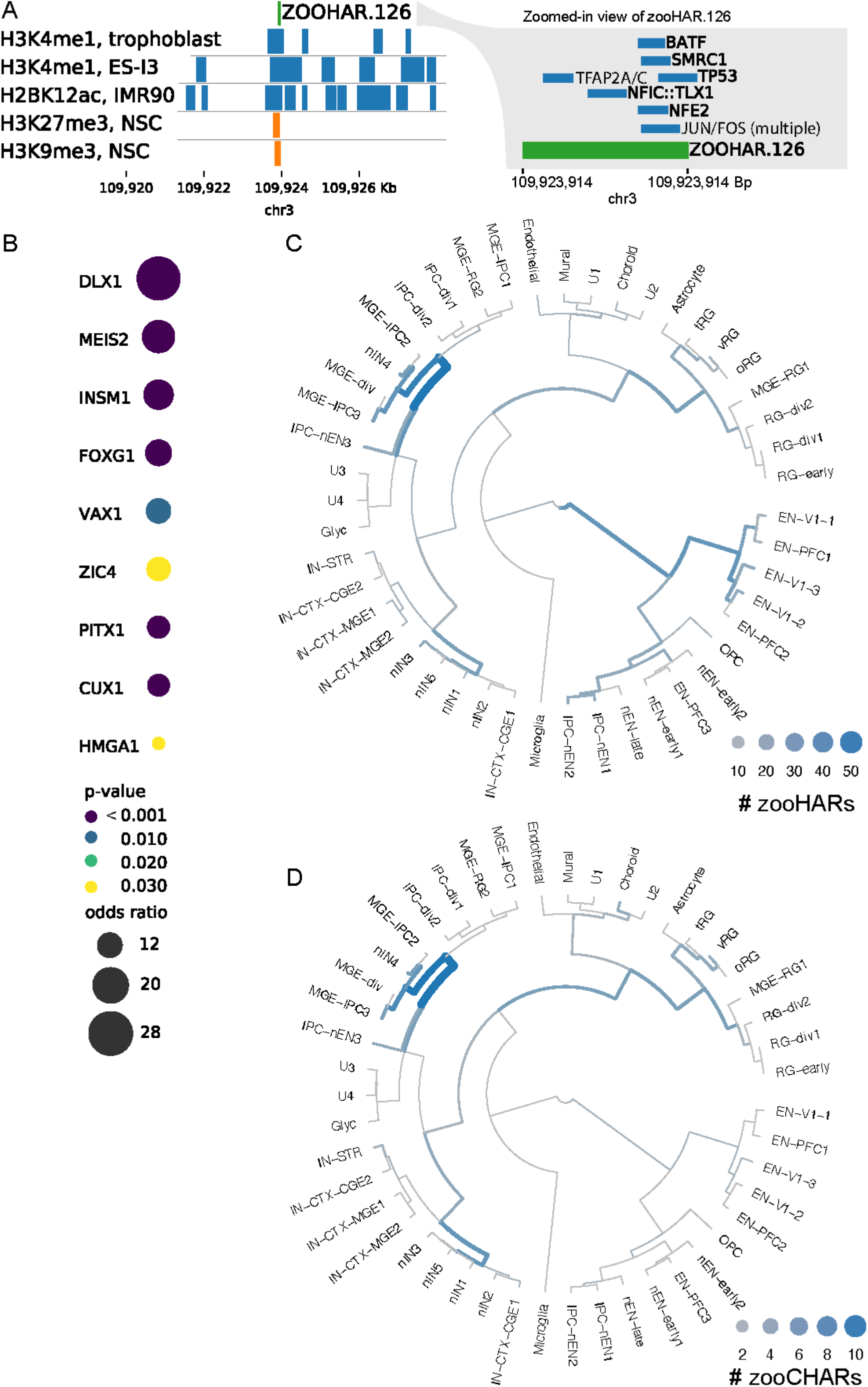
zooHARs in human brain development. (**A**) Transcription factor footprints (*56*) and epigenomic marks (*60*) overlapping zooHAR.126. NSC: neural stem cell. (**B**) Subset of enriched transcription factor footprints in zooHARs relative to phastCons elements (Fisher’s exact p-value ≤ 0.05). Full set available in Table S6. (**C**) Cell types in which zooHARs are predicted to regulate gene expression based on CellWalker analysis of data from the developing human telencephalon. (**D**) Cell type assignments for zooCHARs based on CellWalker analysis of data from the developing human telencephalon. Unlike with HARs, no CHARs map to late stage excitatory neurons. Abbreviations of cell types for (C, D); excitatory neurons (EN) derived from primary visual cortex (V1) or prefrontal cortex (PFC), newborn excitatory neurons (nEN), inhibitory cortical interneurons (IN-CTX) originating in the medial/caudal ganglionic eminence (MGE/CGE), newborn interneurons (nIN), intermediate progenitor cells (IPC), and truncated/ventral/outer radial glia (tRG/vRG/oRG). More cell type information is available at https://cells.ucsc.edu/?ds=cortex-dev (*59*, *65*).

To further specify cell types in the human brain where zooHARs likely function as regulatory elements, we applied the CellWalker method to map them to cell types using single-cell ATAC-seq with RNA-seq from the developing human telencephalon surveyed at mid-gestation (*59*, *65*– *67*). We found the highest number of zooHARs assigned to newborn interneurons, radial glia, excitatory neurons from the prefrontal cortex, and medial ganglionic eminence intermediate progenitors (Fig. 3C, Table S7)(*59*). Repeating this analysis for zooCHARs, cell types were largely similar to those assigned to zooHARs, but many fewer zooCHARs mapped to excitatory neurons from the prefrontal cortex. This difference may provide clues towards the mechanisms underlying species-specific neurodevelopmental traits, such as increased plasticity and protracted maturation in the human brain. However, these results must be interpreted with the caveat that cell-type assignments were made from human data as parallel chimpanzee data are not available (Fig. S9, Table S7). Finally, we repeated the CellWalker analysis using single-cell ATAC-seq and RNA-seq from the human adult brain (*68*, *69*) and heart (*70*). Very few ARs mapped to adult heart cell types. In the adult brain, fewer zooCHARs were assigned cell types compared to zooHARs, with the largest species difference being in excitatory neurons, mirroring our finding in mid-gestation brain (Fig. S10, Table S7).

## Massively parallel validation of zooHARs in human primary cortical cells

To validate these predictions, we performed a massively parallel reporter assay (MPRA) to test the enhancer activity of zooHARs in five replicates of human primary cells from mid-gestation (gestation week 18) telencephalon. Of the 175 zooHARs predicted to function as neurodevelopmental enhancers and passing MPRA quality control, 88 (50.3%) drove reporter gene expression to a level indicative of enhancer activity (Methods; Table S6). This high-confidence set of human accelerated enhancers active in human neurodevelopment includes zooHAR.1, zooHAR.133, zooHAR.138, and zooHAR.156, all of which are in TADs with developmental genes (*GBX2, EFNA5, EN1,* and *PBX3,* respectively) that have differential contacts in our human versus chimpanzee NPC Hi-C data. Prior studies precisely reconstructing human-specific mutations at the endogenous locus in mouse validated zooHAR.1 (also known as HACNS1, HAR2, 2xHAR.3) as an enhancer of *GBX2* and zooHAR.138 (2xHAR.20, HAR19, HAR80) as an enhancer of *EN1.* Other zooHARs with enhancer-like epigenetic signatures but lower MPRA activity may function in different developmental stages or in cell types poorly represented in our telencephalon samples, or their activity may be underestimated by MPRA due to using 270-bp sequences and random integration sites. Despite these limitations, our MPRA data strongly support the conclusion that many zooHARs function as enhancers in cell types of the developing brain. Altogether, this work demonstrates that hsSVs cluster in TADs with HARs that likely function as regulatory elements in neurodevelopment, and these hsSVs can change 3D regulatory interactions of HARs.

## Discussion

Lineage-specific ARs represent sequence-based evolutionary innovations in the genome that may underlie traits that define each species. The Nextflow pipeline introduced in this work enables reproducible identification of ARs in any species in very large alignments, as demonstrated with the Zoonomia dataset of 241 mammals (*23*). Integration of dozens of public and novel datasets refined our understanding of which HARs may function as regulatory elements, at which developmental stages, and in what cell types. Viewing ARs through the lens of 3D genome organization revealed an enrichment of HARs and CHARs in TADs containing species-specific SVs. Generation of the highest resolution cross-species Hi-C dataset to date in matched NPCs from human and chimpanzee enabled further discovery that hsSVs predicted by a deep-learning model to change 3D genome organization nearby HARs and CHARs correspond to true differences between human and chimpanzee NPCs. HARs are active enhancers in diverse cell types and the majority contact putative target genes in a cell type-specific manner (*71*), so future investigation of more cell types may uncover further perturbations.

It is interesting to ask about the sequence of genomic events in loci with hsSVs and HARs. One intriguing possibility is that in some cases the hsSV altered the 3D chromatin contacts of a conserved regulatory element that then underwent rapid adaptation through point mutations in the same species to adjust to its new target genes. With available data, however, we cannot rule out the possibility that the accelerated region changed prior to the structural variant. Nor can we confidently infer that the structural variant and 3D genome changes caused accelerated sequence evolution of the regulatory element. It is also important to note that the vast majority of TADs containing hsSVs with high disruption scores do not contain zooHARs, and about a third contain phastCons elements that are not human-accelerated. Nonetheless, our integrative data analysis points to enhancer hijacking as a potential genetic mechanism to explain HARs and other lineage-accelerated conserved non-coding regions. Further experimentation will be needed to ascertain the validity of this hypothesis. However, it is clear that the evolution of genome sequence and 3D organization do not occur in isolation.

## Materials and Methods

### Automated identification of human and chimpanzee accelerated regions

To facilitate detection of accelerated regions in any lineage on any computing infrastructure, we developed a pipeline implemented in Nextflow (*26*) (Fig. 1A). To date, identification of lineage-specific accelerated regions has used custom scripts that call the PHAST/RPHAST packages (*27*, *29*, *72*) or similar software to identify conserved elements with increased rates of nucleotide substitutions in a given part of a phylogeny using a multiple sequence alignment of the species in the tree. Highly conserved elements are likely to be functional, and they have higher power for detecting accelerated substitution rates on short (e.g., human, chimpanzee) branches as compared to less conserved elements. Our pipeline AcceleratedRegionsNF, available at github.com/keoughkath/AcceleratedRegionsNF, automates these analyses, including tuning run time parameters for large alignments and parallelizing compute over genomic regions (see Supplementary Text). Users provide a multiple sequence alignment in MAF format, a Newick-formatted, bifurcating species tree, and a neutral model. Users may analyze a subset of species in the multiple alignment by also submitting a species list. They simply change the configuration file to describe their computing environment, and the analysis pipeline will run beginning to end. The pipeline generates a BED-formatted file of accelerated regions at a user-defined false discovery rate (FDR) and a table of phastCons elements with phyloP scores and p-values (raw and Benjamini-Hochberg adjusted), enabling the user to adjust the acceleration FDR if desired after running the pipeline. Run time of the pipeline changes based on the size of the computing environment and the size of the input MAF files. Splitting the MAF files into smaller segments (e.g., 10 megabases each) speeds up the runtime significantly.

The human (zooHAR) and chimpanzee (zooCHAR) accelerated regions described in this work were identified using the Zoonomia 241-mammal human-referenced MAF-formatted multiple alignments, a neutral model based on ancestral repeats and the Zoonomia chromosome X species tree (23). Because the multiple alignment was human-referenced, zooHARs and zooCHARs were both initially identified in the human reference genome (hg38). Using the Nextflow pipeline described above, conserved elements in all species in the multiple alignment were identified using phastCons (*72*) with the human or chimpanzee sequence masked, these elements were filtered for level one or two synteny with rhesus macaque, dog, and mouse (*73*). Duplications, pseudogenes from Gencode v29, self-chain and repetitive regions were filtered out (*73*). Elements with a phastCons log odds score in the bottom three deciles were removed, as well as any elements less than 50 base pairs (bp) long. We note that multiple ~100-bp phastCons elements often occur near each other, because a functionally constrained element (e.g., exon, enhancer) may be composed of highly conserved regions broken up by several less conserved alignment columns that cause phastCons to annotate separate conserved regions.

Accelerated elements in human or chimpanzee were identified using phyloP (*28*). Elements with a Benjamini-Hochberg false discovery rate less than 0.05 were retained as accelerated regions. When several phastCons elements are adjacent pieces of a larger enhancer-like element, they were separately tested using phyloP and hence may not all be accelerated.

### Characterization of zooHARs and zooCHARs

zooHAR distribution relative to gene annotations was performed using GENCODE v37 annotation in reference human genome assembly hg38 (*74*). Selection and clustering analyses were conducted as previously described (*10*, *30*). Enriched ontology terms for genes proximal to zooHARs were identified using GREAT (*31*). Functional modules associated with zooHAR-linked genes were detected using HumanBase tissue-specific networks (*32*). Further gene ontology analysis of genes that co-occur in chromatin loops (10kb resolution) with zooHARs was conducted using DAVID (*75*). Epigenomic annotations were performed on the midpoint of each zooHAR extended upstream and downstream by 750bp, and the decision threshold for enhancer predictions adjusted to 0.3, in order to more closely match the properties of validated VISTA enhancers (*64*). zooHAR brain cell types were identified by CellWalker (*65*) as implemented in the CellWalkR package (version 0.99, default parameters, with Jaccard similarity used for cell edges, gene accessibility used for label edges, and the label edge weight parameter set to one)(*66*) applied to data from the developing human telencephalon (*59*, *65*). zooHAR expression was assessed by overlap with transcripts from (*76*) lifted over to hg38 (*77*). Enrichment of zooHARs in chromatin contact domains with human-specific structural variants (hsSVs) was performed by calculating the odds ratio of a chromatin contact domain containing a zooHAR and an hsSV. A p-value was generated by comparing that odds ratio to a null distribution of 1000 odds ratios calculated the same way, except with a random draw of N phastCons elements, where N is the number of zooHARs. Various computational analyses utilized GNU parallel (*78*). To characterize chimpanzee accelerated regions, the above analyses were repeated with zooCHARs in place of zooHARs.

### Prediction *in silico* of human-specific structural variant impacts

Prediction of hsSV effects was performed using Akita, a deep learning model that predicts chromatin contact matrices from DNA sequence (*36*). To predict the impact of hsSVs on the 3D genome, we submitted two 1Mb sequences to Akita, one with and one without the hsSV. We used the human (hg38) sequence if the hsSV was an insertion and chimpanzee (pantro6) sequence if the hsSV was a deletion or inversion. We then calculated the mean squared error (“disruption score”) between these two contact matrices.

### NPC generation, differentiation, validation

Two human (WTC11 and HS1) and two chimpanzee (C3649 and Pt2a) induced pluripotent cell lines (iPSCs) were cultured in Matrigel-coated plates with mTeSR media (WTC11 and C3649 were cultured in StemFlex) in an undifferentiated state. Cells were propagated at a 1:3 ratio by treatment with 200 U/mL collagenase IV (or PBS-EDTA) and mechanical dissection.

WTC11 and C3649 iPSCs were differentiated to neural progenitor cells (NPCs) and validated as previously described (*37*). Briefly, 2-2.5×10⁵ cells per cm² were seeded on Matrigel-coated wells in StemFlex containing 2 μM Thiazovivin. The following day (Day 0), medium was replaced with E6 containing 500 nM LDN193189 (Selleckchem), 10 μM SB431542 (Selleckchem), and 5 μM XAV-939 (Selleckchem). Starting on Day 3, medium was replaced with E6 containing 500 nM LDN193189 and 10 μM SB431542 every 48 hrs. Starting on Day 12, medium was replaced with Neurobasal containing 2 mM GlutaMAX, 60 μg per ml L-Ascorbic acid 2-phosphate, N2, and B27 without Vitamin A every 48 hours. Around Day 16, cells were washed with PBS, dissociated with Accutase, pelleted and resuspended in Neurobasal containing 2 mM GlutaMAX, 60 μg per ml L-Ascorbic acid 2-phosphate, N2, and B27 without Vitamin A, 10 ng per ml fibroblast growth factor 2, and 10 ng per ml epidermal growth factor, and seeded on poly-L-ornithine-, fibronectin-, and laminin-coated wells. Cells were collected for HiC at passage 5-7.

To differentiate HS1 and Pt2a iPSCs into NPCs, cells were split with EDTA at 1:5 ratios in culture dishes coated with matrigel and culture in N2B27 medium (comprised of DMEM/F12 medium (Invitrogen) supplemented with 1% MEM-nonessential amino acids (Invitrogen), 1 mM L-glutamine, 1% penicillin-streptomycin, 50 ng/mL bFGF (FGF-2) (Millipore), 1x N2 supplement, and 1 x B27 supplement without Vitamin A (Invitrogen)) supplemented with 100 ng/ml mouse recombinant Noggin (R&D systems). Cells at passages 1-3 were split by collagenase into small clumps, and continuously cultured in N2B27 medium with Noggin. After passage 3, cells were plated at the density of 5×10⁵ cells/cm² after disassociation by TrypLE express (Invitrogen) into single-cell suspension, and cultured in N2B27 medium supplemented with 20 ng/mL bFGF and EGF. Cells were maintained and collected at passage 18-20. Our use of two differentiation protocols reflects rapid progress in stem cell research during the course of this study. Cells from the same populations were validated and used in a previous study (*37*). We verified that the chromatin interactions in the resulting Hi-C data did not show a batch effect across protocols.

### Hi-C data generation

Hi-C was performed using the Arima Hi-C kit (Arima Genomics) according to the manufacturer’s instructions. 10 million cells were used. The sequencing library was prepared using Accel-NGS 2S Plus DNA Library Kit (Swift Biosciences) according to the manufacturer's protocol. Two independent biological replicates were prepared for each cell line. In total eight libraries were pooled and sequenced with paired-end 150-bp reads using two lanes of a NovaSeq6000 S2 (Illumina) at the Chan Zuckerberg Biohub.

### Hi-C data processing

Adapters were trimmed from raw FASTQ files using TrimGalore [v0.6.5] with options -- illumina --paired. The data were then processed from adapter-trimmed FASTQ files to Hi-C contacts as cooler files using Distiller [v0.3.3] (*79*). This processing includes read mapping with BWA-MEM (*80*), filtering (MAPQ >= 30), contact pair processing with pairtools (*81*) and normalization via matrix balancing (*82*). Samples were processed both per replicate, per individual and per species. For easier comparison of samples in some analyses, we mapped the data from each species to the reference genome of the other species (human to pantro6 and chimp to hg38). Cis/trans ratio was calculated as the ratio of cis to trans contacts for each replicate (*83*). Distance-dependent interaction frequency decay was computed using cooltools with 100-kilobase (kb) bins (*83*, *84*).

A and B compartments were identified by eigenvector decomposition of the contact matrices, phased by GC content with A compartment having higher GC content than B compartment using cooltools (*79*). We assessed conservation between TAD boundaries based on the method from (*42*). We identified boundaries by calculating the insulation score at a resolution of 50kb and using a 800-kb sliding window, considering bins with boundary strength greater than 0.1 and insulation score less than zero as boundaries. Boundaries were considered conserved if they were within two bins (100 kb) of a boundary in the other species, and species-specific if they were more than five bins (500 kb) from the nearest boundary in the other species after liftOver (*42*). TADs for the zooHAR enrichment analyses were identified using a 400-kb window and 10-kb bin size, with boundary strength greater than 0.1 and insulation score less than zero as boundaries. Loops were identified using Mustache at 5-kb resolution (*46*). Conservation of loop anchors was conducted using mapLoopLoci (*47*).

### Massively parallel reporter assay

We designed 270-bp oligos centered on zooHARs and positive control enhancer sequences. For zooHARs longer than 270 bp, we tiled oligos across the element. A 31-bp minimal promoter (minP) and 15-bp random barcodes were placed downstream of the synthesized oligos via PCR and cloned into an MPRA vector as previously described (*85*). The library was packaged into lentivirus and used to infect human primary cortical cells dissociated from two fresh tissue samples (gestational week 18). Cells were cultured for two days prior to infection and 3 days following infection in a DMEM-based media containing B27, N2, and Pen-Strep. Cells were harvested, then DNA and RNA were obtained for sequencing. For each oligo, we quantified enhancer activity using the ratio of barcode abundance in RNA versus DNA normalized and batch corrected across replicates. A zooHAR was determined to be active if its maximally active tile had an RNA/DNA value exceeding the median of a set of positive control enhancer sequences that we included in the MPRA library.

## Supplementary Text

### Impact of number and choice of species in the alignment

Previous analyses to identify human accelerated regions (HARs) have generally used alignments of fewer than 30 species (*1*, *2*, *51*). The Zoonomia multiple alignment analyzed in this work, as well as other commonly used multiple alignments of vertebrates, such as the 100-way UCSC alignment, are much larger. Additionally, genome quality and completeness for many species have improved greatly since early HAR analyses. Therefore, we systematically assessed each step of the HAR identification analysis laid out in earlier work to determine whether changes needed to be made.

In order to assess the variability of HARs and phastCons elements per species number and set, we identified HARs and phastCons elements from the 100-way hg38 UCSC multiple alignment of vertebrates using sets of ten to ninety randomly selected species with three replicates of random species selection per species number. Each species set included human and chimpanzee, but was otherwise randomly selected from the full set of species in the UCSC 100-way alignment. These HARs were identified using a neutral model based on 4-fold degenerate sites, phastCons parameters rho=0.3, omega=45, gamma=0.3 with a Benjamini-Hochberg FDR < 0.01 for phyloP acceleration. We found that with increasing numbers of species, the number of elements identified, genome coverage, and size of elements all decreased (Fig. S11). These trends were consistent across other FDR thresholds. We next compared the HARs analyzed in the main text using the Zoonomia 241-mammal alignment (zooHARs) to HARs identified from the subset of all mammals (UCSC mammal) and from the full set of species (UCSC vertebrate) in the hg38 100-way MULTIZ alignment from UCSC, in each case based on a neutral model derived from ancestral repeats. Most UCSC vertebrate HARs were a subset of the UCSC mammal HARs or zooHARs (Fig. S12), while UCSC mammal HARs and zooHARs shared about half of their elements and base pairs. These results indicate that the alignments used to identify phastCons elements have a big impact on the resulting set of ARs, and including non-mammal vertebrates decreases the number of ARs discovered.

### Using a subset of high-quality species for HAR identification

We explored the strategy of using a subset of “high-quality” species genomes for HAR identification, with the rationale that this may help avoid false positives caused by spurious alignments or miscalled regions in genomes. A barrier to this approach was that genomes from different species were assembled using different sequencing technologies and methodologies, making it difficult to establish a set of objective standards for inclusion. Additionally, many of the “higher quality” genomes are in the primate clade, thus skewing the phylogenetic representation of the species set. Due to these constraints, we were not able to curate an optimal species set based on maximizing stability of the HARs identified. Therefore, we decided to proceed with the full set of species to identify zooHARs. However, these results emphasize the importance of careful species set selection in AR analyses depending on the research goals. To this end, in the AR-identification pipeline described in this paper, we enable the researcher to submit a list of species in order to analyze a subset of the species present in the multiple sequence alignment.

### Tuning phastCons parameters in assemblies with hundreds of species

The methods to identify HARs were developed using alignments with less than 30 species and older versions of genome assemblies. As multiple species alignments have grown and assemblies have improved and become more complete, we systematically assessed the parameter choice for identifying the set of conserved elements from which HARs would be drawn. The tuning parameters in phastCons that we assessed included ρ, a scaling factor describing the extent to which a neutral tree should be shrunk to approximate the conserved state, ω, the estimated length of conserved elements, and ɣ, the estimated genome coverage by conserved elements. Of these parameters, the most obvious candidate to be adjusted was ɣ, as this parameter is inversely proportional to the proportion of the reference genome in the multiple alignment blocks. In previous alignments, only 16.5% of the human genome was represented in alignment blocks, whereas in the Zoonomia 241-mammal alignment that has increased to 97.7%. Therefore, based on (*72*) and an expected genome coverage of 5% by conserved elements, ɣ is approximately 0.05. As another method of checking these parameters, we estimated ρ, ω and ɣ by maximum likelihood using the phastCons program. The parameters were estimated based on 100 1-Mb windows of the UCSC 100-way alignment, using a neutral model estimated from ancestral repeats. The median values identified were ɣ=0.06, ρ=0.27 and ω=4.05. Thus, we decided to proceed with parameters ɣ=0.05 and ρ=0.3 based on these estimates, but we used ω=45 as done in previous work with the goal of increasing the size of the conserved elements identified, which increases power in downstream phyloP tests for acceleration (27) and eliminates the need to develop ad hoc methods to merge adjacent phastCons elements. Additionally, we implemented a threshold for the phastCons log odds score, requiring that phastCons elements considered for acceleration were above the third decile of length-normalized log odds scores, thus removing elements with the weakest signatures of conservation from consideration.

### Automated identification of human- and chimpanzee-specific accelerated regions

Genome-wide analyses of large multiple-species alignments typically require cluster computing, which hinders reproducibility and accessibility. To enable automated detection of accelerated regions in any lineage on any computing infrastructure, we implemented our analysis pipeline in Nextflow (*26*) Given a species tree, neutral model, and multiple sequence alignment, this open-source software uses PHAST to identify lineage-specific accelerated regions for any species of interest (Fig. S1A). This pipeline enables simplified, portable and reproducible identification of lineage-specific accelerated regions.

### zooHAR and zooCHAR characterization

#### Accelerated regions cluster and are mostly noncoding

Using the Zoonomia 241-mammal alignments, we identified 312 zooHARs and 141 chimpanzee accelerated regions (zooCHARs) (Benjamini-Hochberg FDR < 0.05, Tables S1, 2). Median length was 117.5 base pairs (bp) for zooHARs (IQR: 110.5 bp) and 108.0 bp for zooCHARs (IQR: 90 bp), similar to prior studies. 32.4% of zooHARs overlap previous lists of HARs identified by similar methods (*1*, *6*, *51*), and 5.5% of a merged group of previous sets of HARs identified by varying methods (*1*, *3–5*, *9*, *51*) overlap zooHARs, agreeing with prior analyses which found that differing methodologies and underlying datasets render most HAR sets unique from one another, and thus we do not claim this set to be superior to others (*9*). zooHARs and zooCHARs were identified on all autosomes and chromosome X. Each set is clustered along the linear genome so that specific loci harbor more zooHARs (p = 0.01) or more zooCHARs (p = 0.01) than expected given the density of conserved (phastCons) regions (225,317 phastCons elements from which zooHARs and 225,287 from which zooCHARs were identified). zooHARs and zooCHARs show a similar genomic distribution to previous HAR sets with respect to genomic features. The majority are intergenic, although some overlap protein-coding features and noncoding RNAs (Fig. S1B, C).

Genes near zooHARs are involved in transcriptional regulation, forebrain development and morphogenesis, and multiple other developmental terms based on GREAT analysis (Fig. S2A) (*31*). GREAT analysis also revealed enrichment of zooHARs nearby genes involved in mouse neonatal lethality with complete penetrance, and multiple abnormal developmental events in mouse (Fig. S2B). GREAT analysis of zooCHARs revealed an enrichment of nearby genes for transcriptional regulation and sequence specific DNA binding (Fig. S3) and neonatal lethality (*31*). However, GREAT analyses are based on genes nearest HARs and CHARs, which may not be the target genes of these elements. Therefore we also identified ontology terms enriched in genes that are associated with HARs and CHARs via 3D chromatin loops from the Hi-C data in NPCs generated in this study. Enriched gene ontology (GO) terms included multiple developmental terms, including “heart development”, “positive regulation of developmental process”. Thus, regardless of the method for associating zooHARs and zooCHARs with target genes, we see a clear enrichment for loci with transcription factors in both species. Developmental processes are also enriched, particularly for zooHARs. The stronger signal for diverse developmental processes in zooHAR loci as compared to zooCHAR loci may be due to higher power with the larger set of zooHARs, but it could also reflect biological differences in the function of these elements in the two species, consistent with adaptation of each species to their distinct environmental niches.

#### Most zooHARs and zooCHARs are under positive selection

Accelerated evolution is not synonymous with positive selection. Positive selection indicates a rate of nucleotide substitutions that is faster than the (local or genome-wide) neutral rate, indicating that the sequence changes are beneficial. Acceleration means a rate of nucleotide substitutions that is faster than expected given the rate in the rest of the tree, which could be faster, slower or equal to the neutral rate. The rest of the tree is evolving slower than the neutral rate for the accelerated regions in this study, so the lineage of interest (human or chimpanzee) could be less slow but still below the neutral rate, equal to the neutral rate or faster than the neutral rate. GC-biased gene conversion (GBGC) can mimic positive selection (*86*), but the substitutions are biased towards A/T to G/C changes. To infer the evolutionary forces that shaped the accelerated regions in this study, we applied a method that uses likelihood ratio tests to assess loci for evidence of positive selection, GBGC, or both (*10*). This method controls for local variation in the neutral rate of evolution by comparing each element to the surrounding 1 Mb of genome rather than the rate of evolution in the other species without the element itself (based on rescaling a phylogeny built using the genome-wide neutral rate). This analysis estimated that 82% of zooHARs and 86% of zooCHARs are under positive selection, though 7% of zooHARs and zooCHARs show strong evidence for GBGC, and 5% of zooHARs may have been shaped by a combination of selection and GBGC (Fig. 1D, E; Tables S1, S2).

#### zooHARs and zooCHARs are transcribed in the developing human brain

Some HARs have been shown to function as noncoding RNAs, including the original HAR1 (*2*), therefore we investigated the noncoding RNA potential of zooHARs. Additionally, many active enhancers are transcribed (eRNAs). We assessed the expression of zooHARs and zooCHARs in RNAseq data from the developing human neocortex (*76*), including both poly-A and total RNA, enabling the study of non-protein-coding RNA transcripts (*76*) and eRNAs. We found that 100 of 312 zooHARs (32%) and 41 of 141 zooCHARs (29%) were expressed in the total RNA dataset (TPM>5, Fig. S1E, F). Twenty of the expressed zooHARs overlapped gene exons, including *ERC2*, involved in neurotransmitter release (*87*), and *TNIK*, implicated in neurological disorders, neurogenesis and cell proliferation (*88*). Of the expressed zooHARs 88 overlapped gene introns, 12 overlapped annotated noncoding RNAs, and 13 do not overlap any currently annotated elements, and therefore could represent uncharacterized noncoding RNAs or eRNAs.

**Fig. S1.**
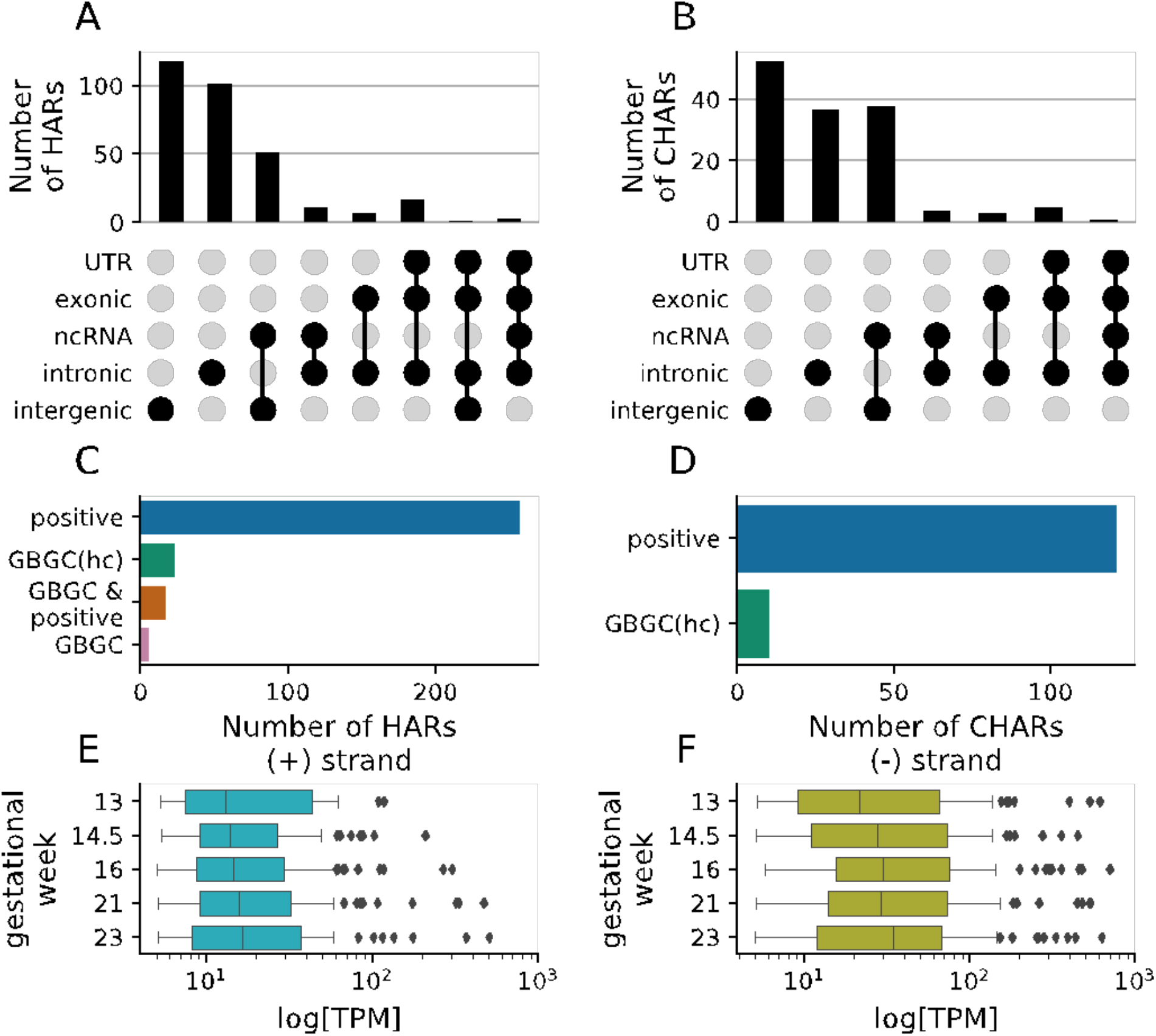
zooHARs demonstrate similar characteristics to prior HAR sets. (**A, B**) Genic distribution of zooHARs (A) and zooCHARs (B) (both in reference hg38), based on Gencode V37 annotations. (**C, D**) Selective forces acting on zooHARs (C) and zooCHARs. (D) from pipeline described in (*10*). Positive=positive selection, GBGC=GC-biased gene conversion, hc=high-confidence. (**E, F**) Transcription of zooHARs from the positive (E) and negative (F) strand in the developing human neocortex at five mid-gestation time points (*76*). Whiskers extend to 1.5 times the inter-quartile range (IQR). TPM=transcripts per million.

**Fig. S2.**
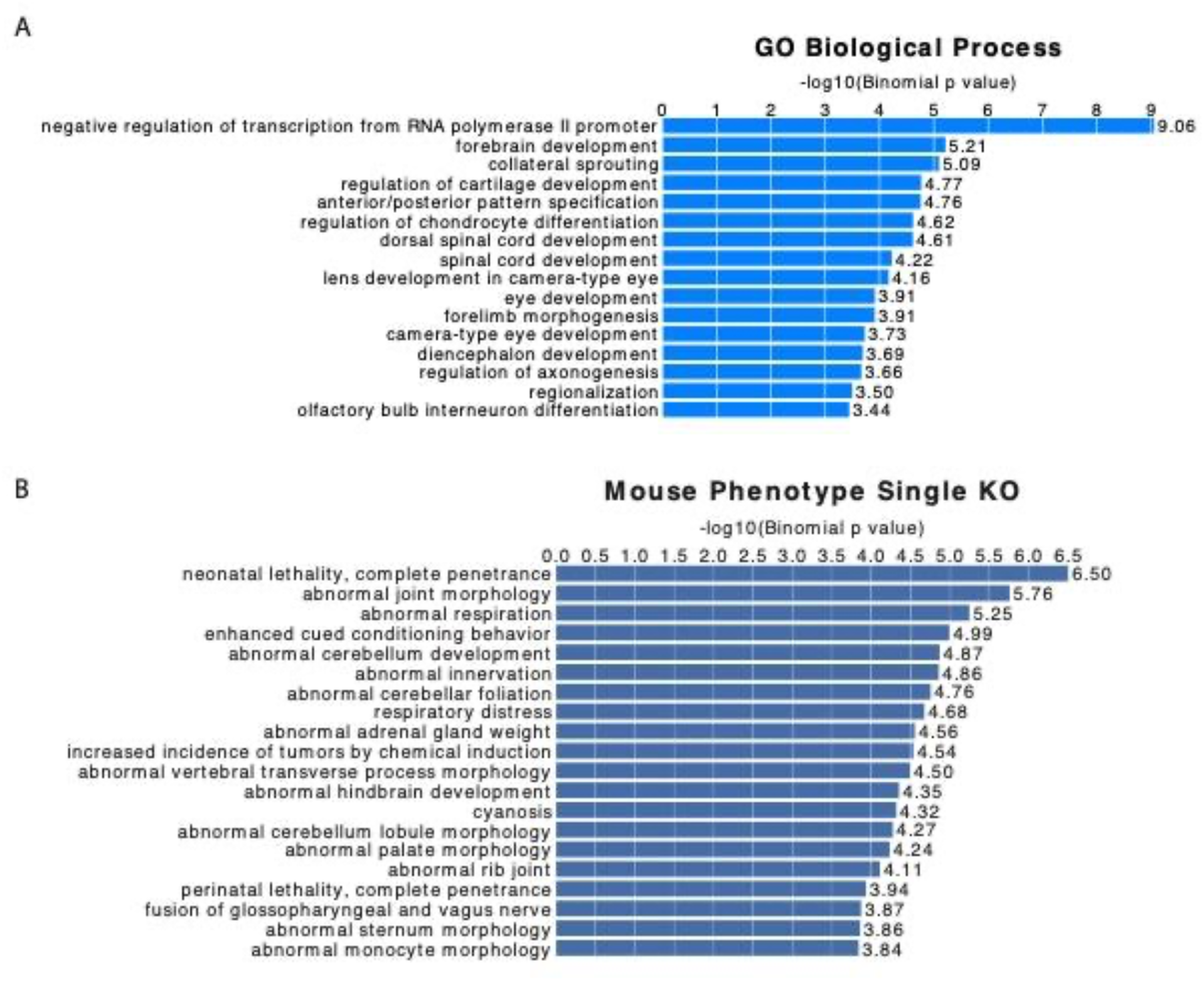
GREAT analysis of genes near zooHARs. (**A**) GREAT (*31*) gene ontology enrichment analysis of zooHARs. (**B**) GREAT mouse phenotype (single knockout) enrichment analysis of zooHARs.

**Fig. S3.**
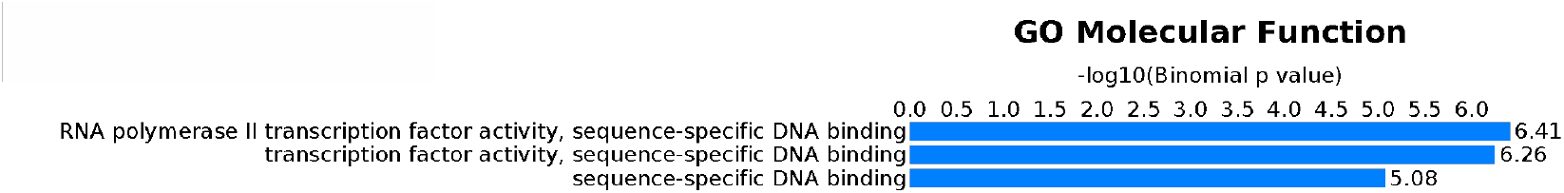
GREAT analysis of genes near zooCHARs. GREAT (*31*) gene ontology analysis of zooCHARs.

**Fig. S4.**
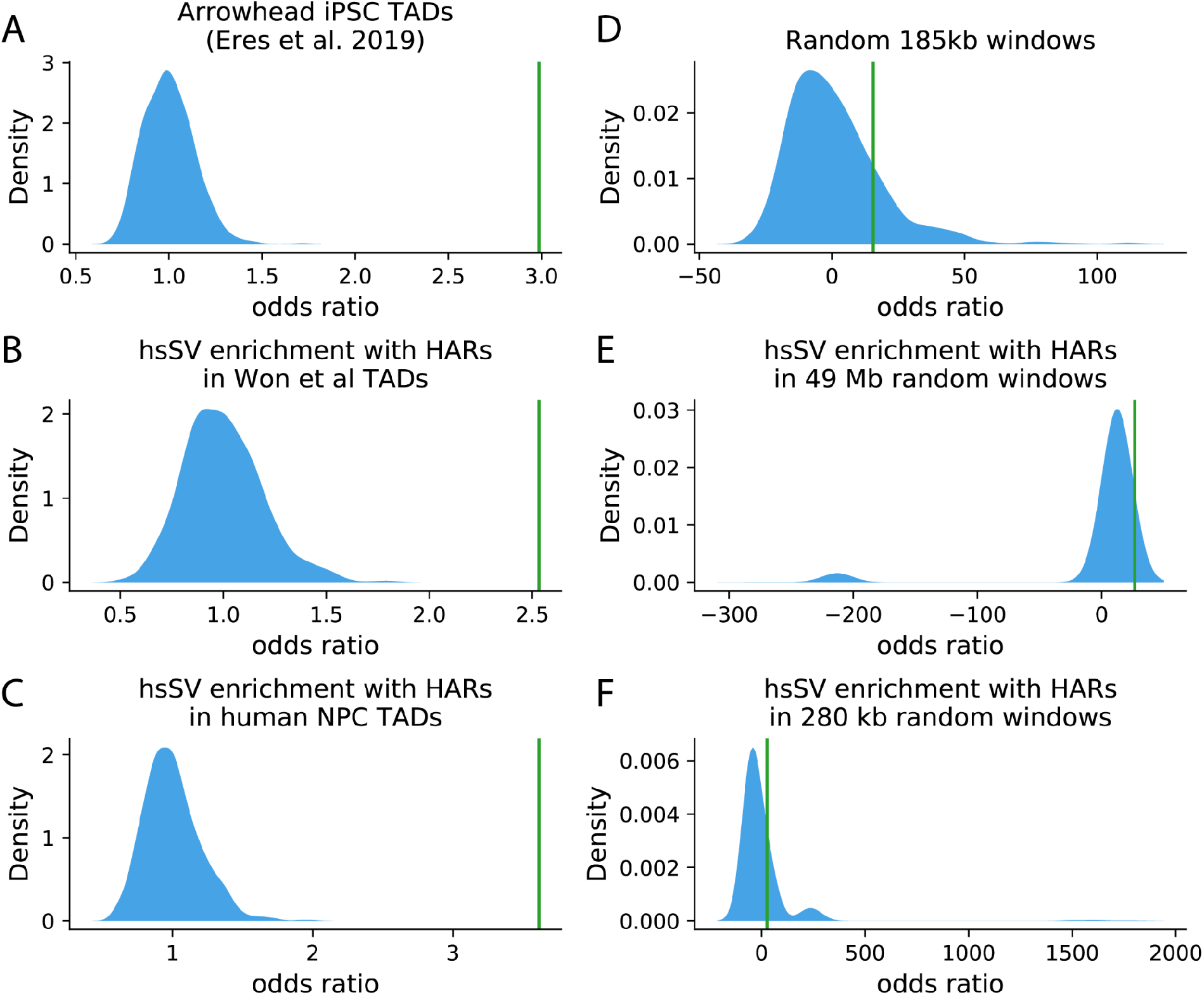
Enrichment of zooHARs in TADs with hsSVs compared to random windows. (A) Odds ratio of TADs from human iPSCs called with the Arrowhead algorithm (*41*) containing one of the 17,789 human-specific structural variants (hsSVs) and one of the 312 zooHARs (green line) compared to a null distribution based on 1000 random draws of 312 phastCons elements (blue shaded area). (B-F) Same analysis as in A, but with (B) TADs from mid-gestation developing human cerebral cortex (cortical plate and germinal zone) based on insulation scores (*89*); (C) TADs from human NPCs (*25*); (D) random 185-kb windows, the median size of contact domains from (A); (E) random 185-kb windows, the median size of contact domains from (B); (F) 280-kb random windows, the median size of the human NPC TADs from (C).

**Fig. S5.**
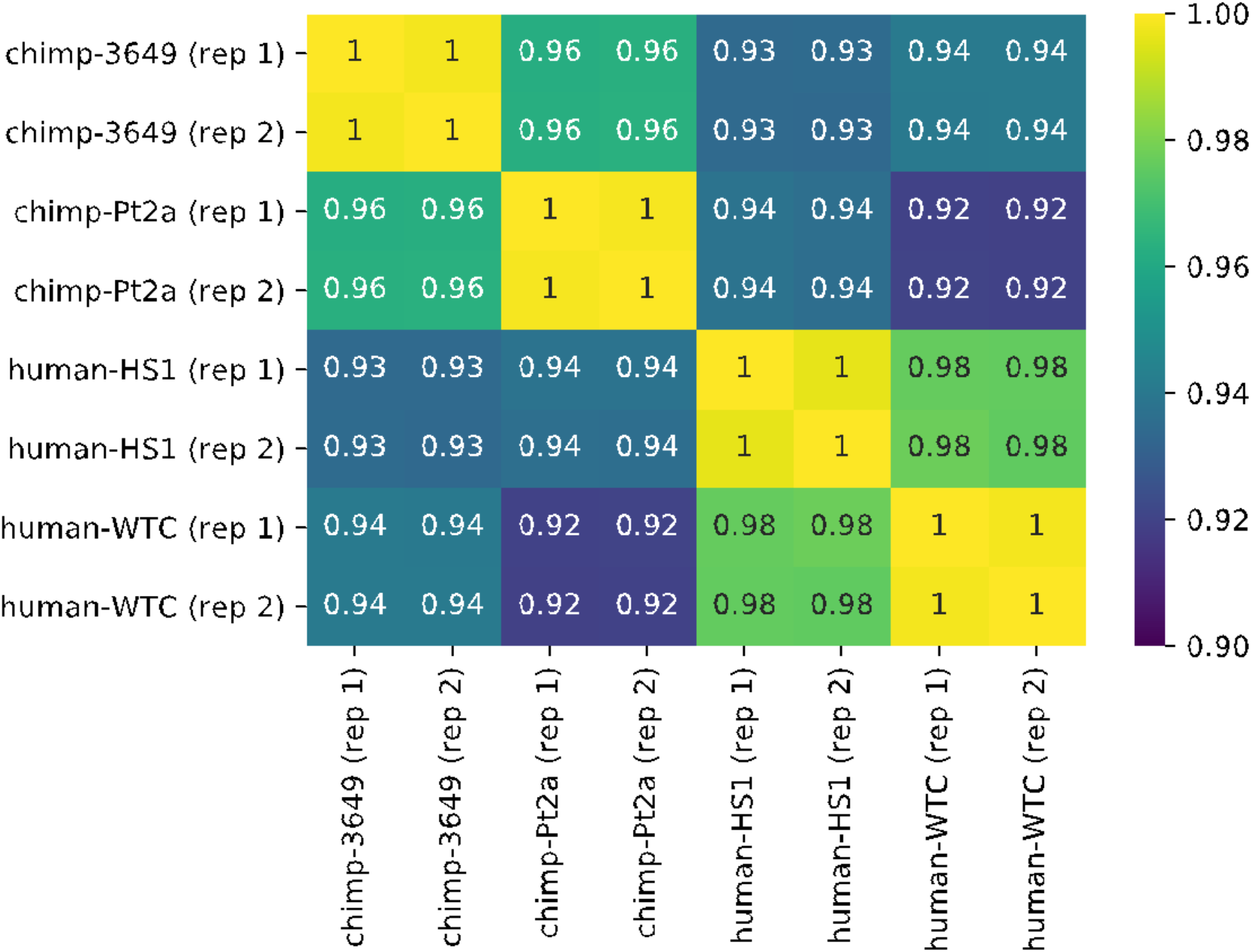
Hi-C correlation values per Hi-C sample. Stratum adjusted correlation coefficients (SCC) between all samples mapped to hg38. The SCC statistic is calculated by stratifying the data by genomic distance, then computing a Pearson correlation coefficient for each stratum and then aggregating the stratum-specific correlation coefficients using a weighted average, with the weights derived from the generalized Cochran– Mantel–Haenszel statistic (*38*).

**Fig. S6.**
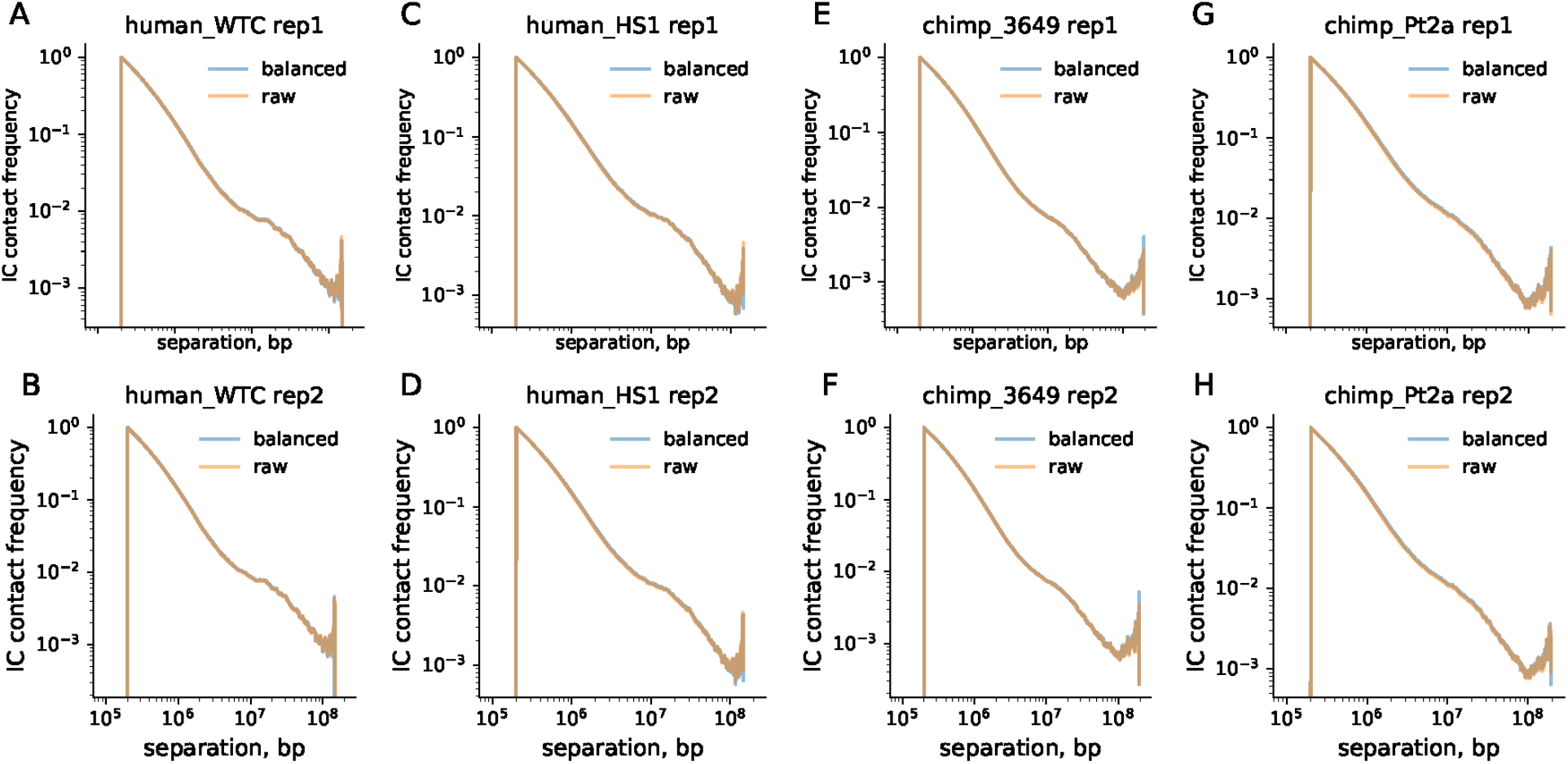
Distance-dependent contact decay per Hi-C sample. Corrected (IC) contact frequency as a function of distance between all pairs of 100-kb bins for each replicate.

**Fig. S7.**
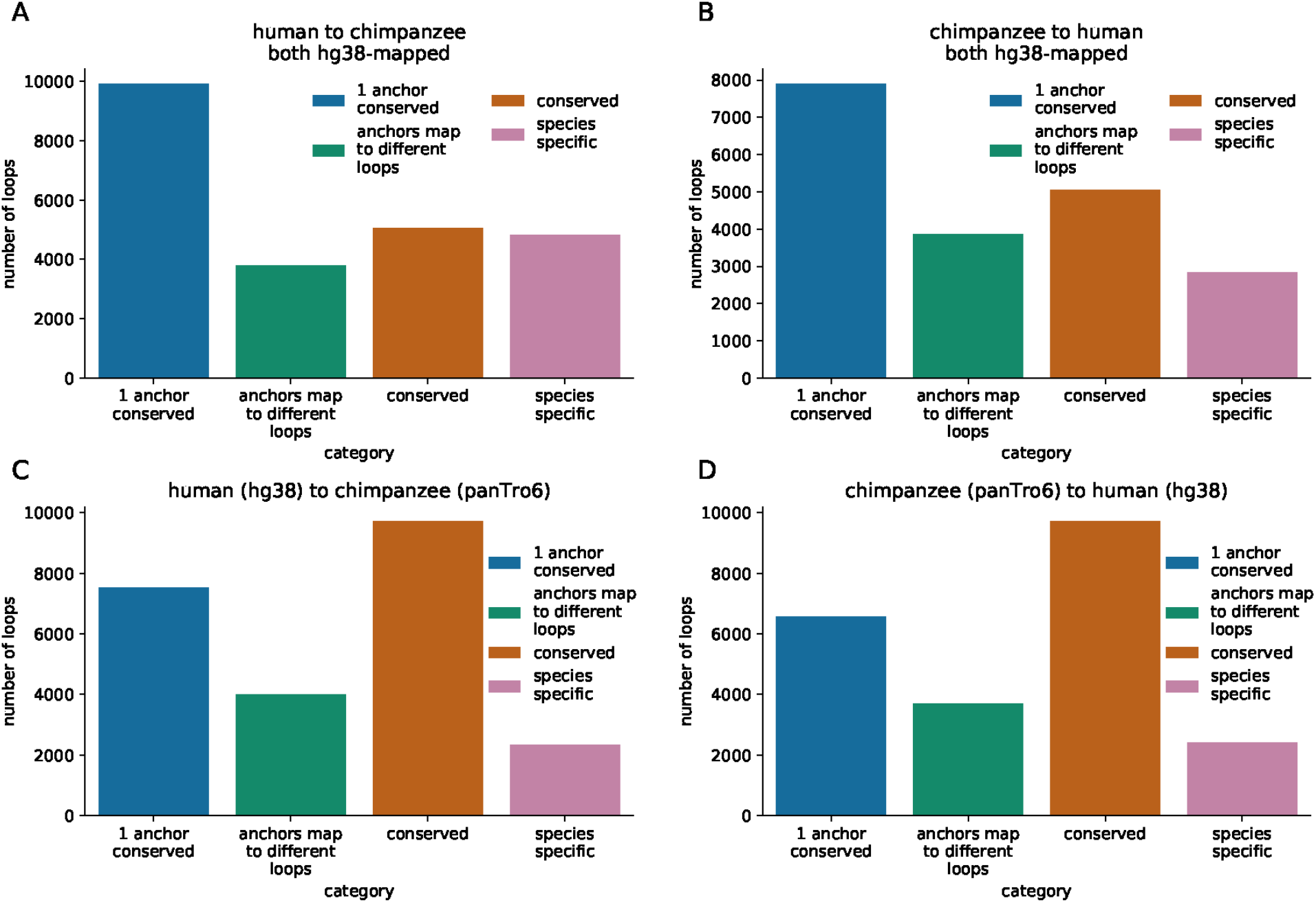
Loop conservation human to chimpanzee. Loop conservation assessed with mapLoopLoci from (*47*) for human compared to chimpanzee Hi-C data with (**A, B**) both mapped to hg38 and (**C, D**) each mapped to their respective species’ reference genome.

**Fig. S8.**
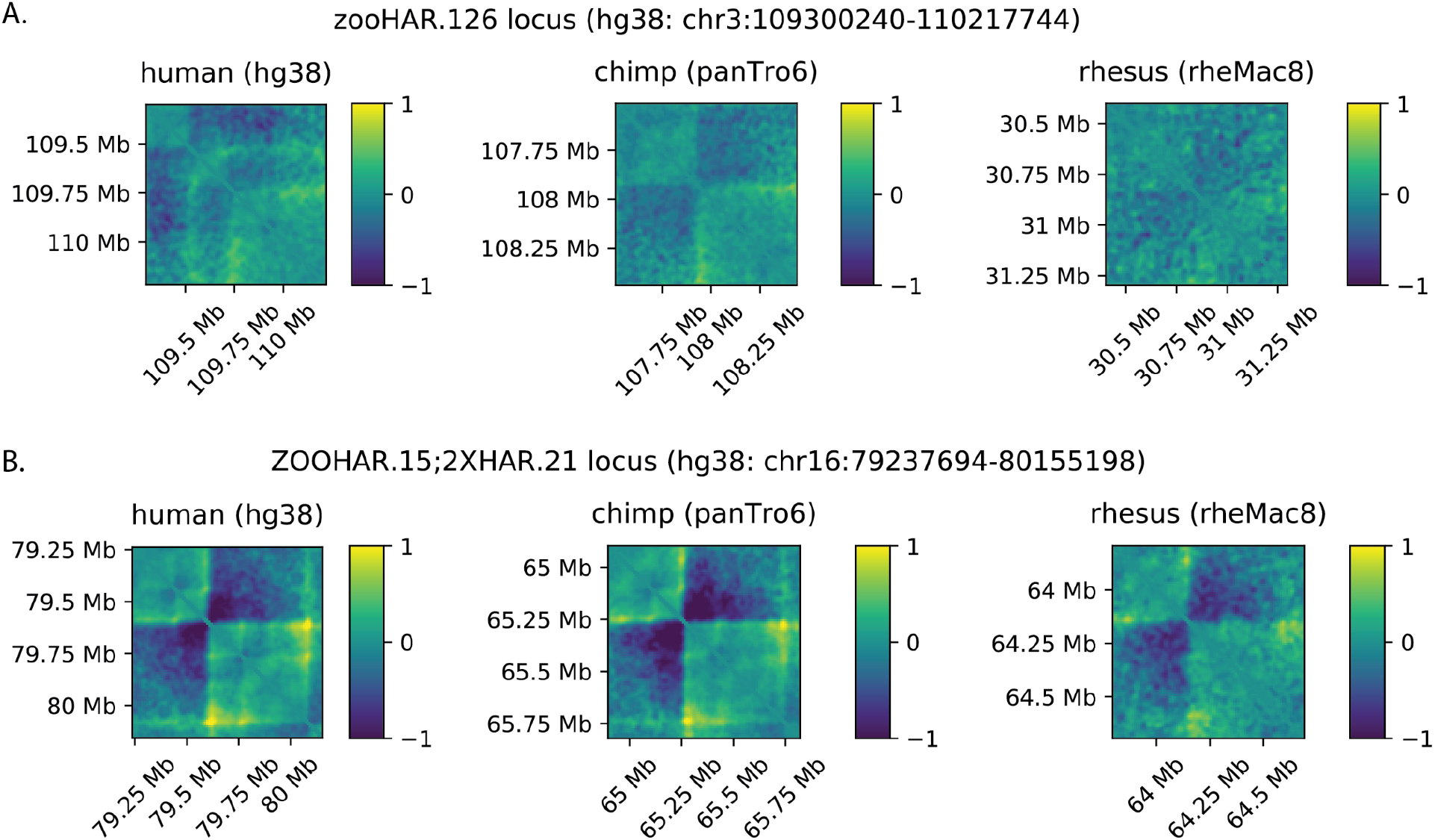
Loci of interest in rhesus macaque. The loci surrounding zooHAR.126 (**A**) and zooHAR.15 (**B**) in human and chimpanzee NPC Hi-C generated in this work, compared to Hi-C from rhesus macaque cortex plate (*42*). Log(observed/expected) values are shown in the heatmaps.

**Fig. S9.**
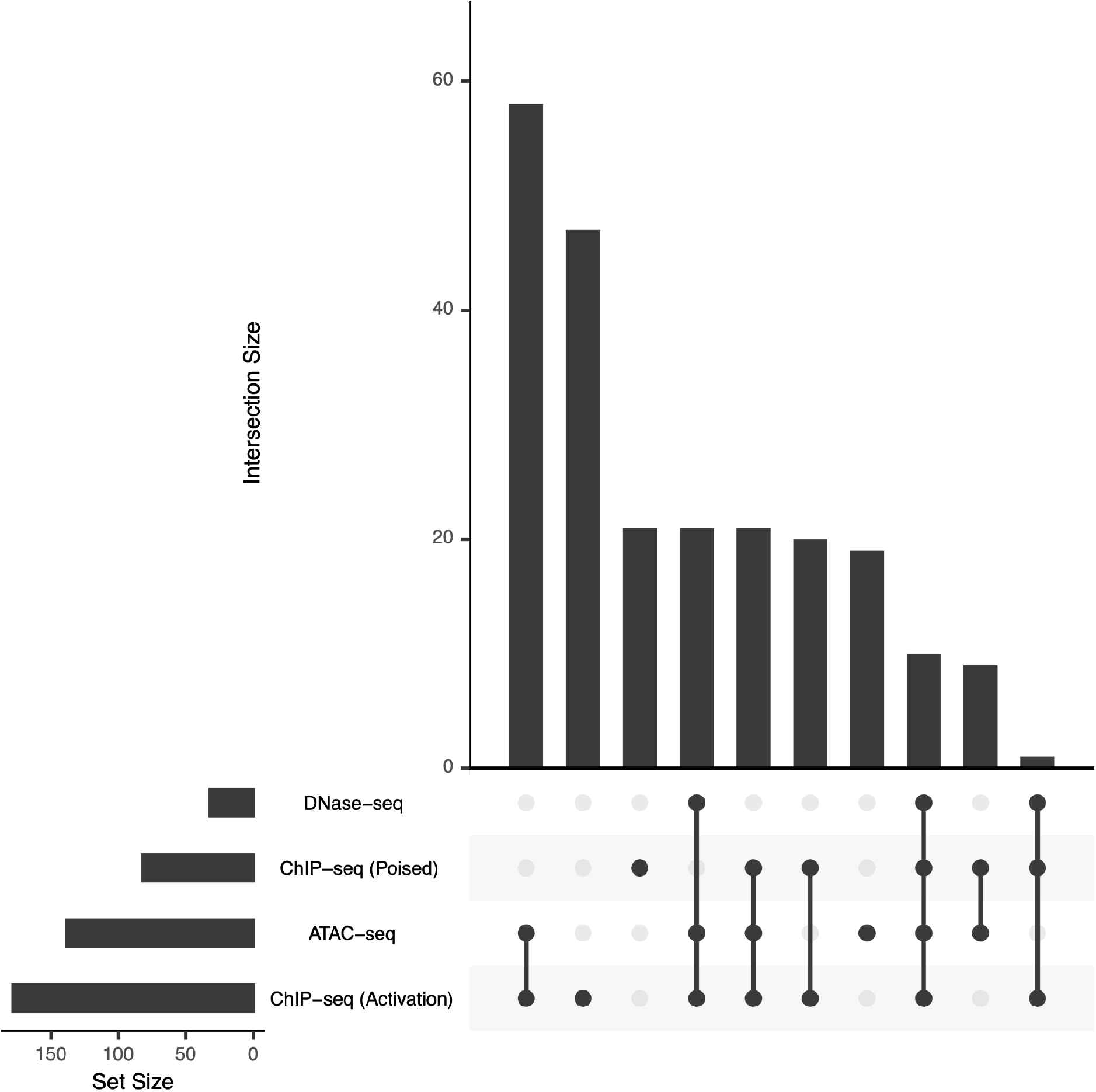
Overlap of zooHARs with epigenomic marks from brain. A majority of zooHARs overlap robust peaks (Irreproducible Discovery Rate 10%) from open chromatin (ATAC-seq or DNase-seq) and activating histone modifications (ChIP-seq) from neural cell lines or primary brain tissue (*53–60*). Bar plot on the y-axis (left) indicates the number of zooHARs overlapping each epigenomic feature, bar plot on the x-axis (top) indicates the number of zooHARs overlapping multiple epigenomic features, indicated by the shaded dots in the center. The highest proportion of zooHARs overlap both activating ChIP-seq and ATAC-seq peaks, followed by those that overlap only activating ChIP-seq peaks. Accounting for the smaller number of DNase-seq datasets, many zooHARs that overlap activating ChIP-seq and ATAC-seq also overlap DNase-seq peaks.

**Fig. S10.**
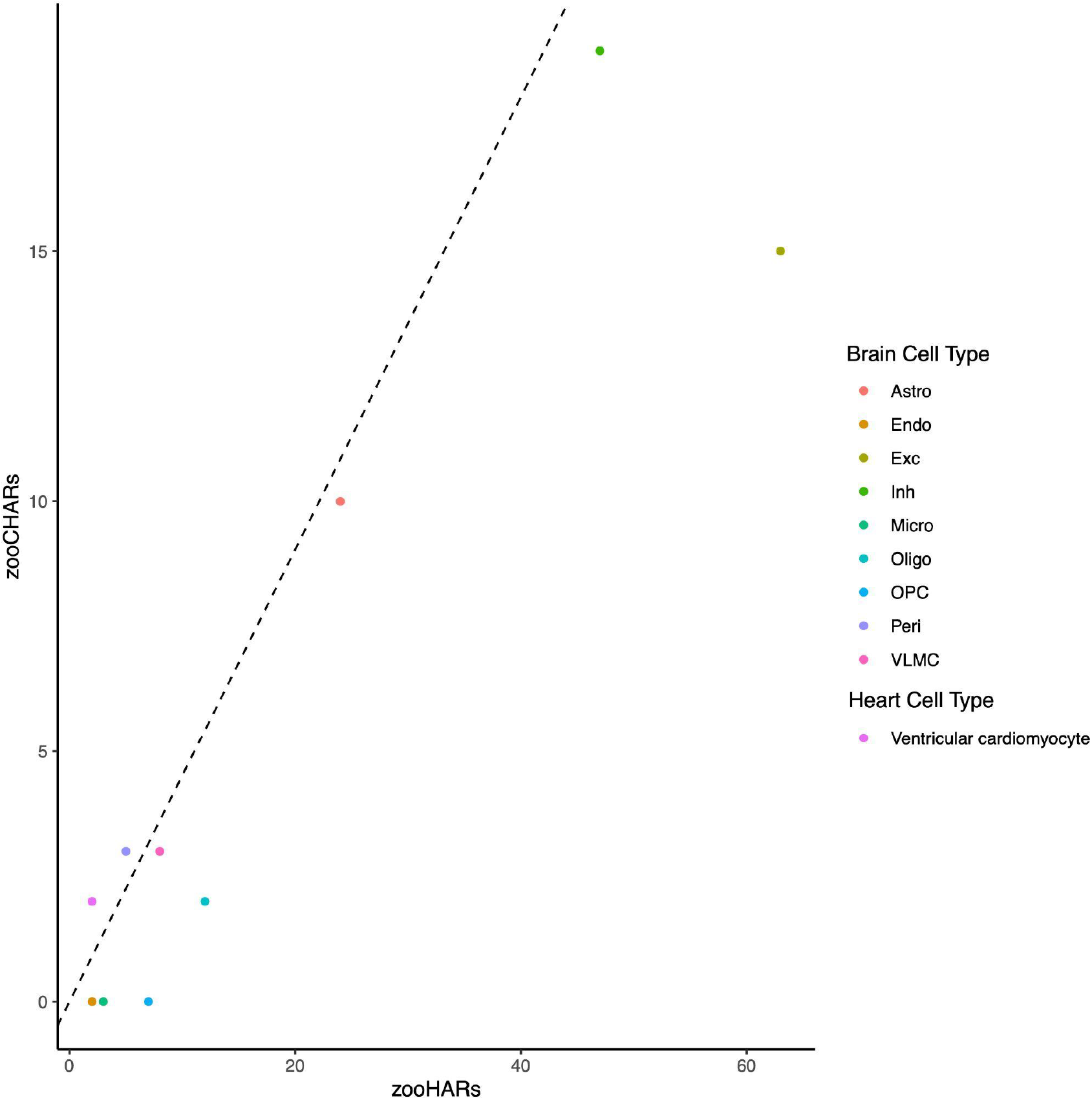
CellWalker analysis of zooHARs and zooCHARs mapped to adult brain and heart cell types. As controls to compare to our analysis of mid-gestation telencephalon cell types, we ran CellWalker using matched single-cell ATAC-seq and RNA-seq from adult brain (*68*, *69*) and adult heart (*70*) to associate each zooHAR and each zooCHAR with cell types in which they appear to be active. The only heart cell type with any ARs is ventricular cardiomyocytes, which was predicted as an active cell type for only a few zooHARs and zooCHARs. In both adult tissues, cell types tend to have similar numbers of zooHARs and zooCHAR associations, with the exception of excitatory neurons, which have many more zooHAR associations. This enrichment mirrors what we observed in mid-gestation brain excitatory neurons (Fig. 3).

**Fig. S11.**
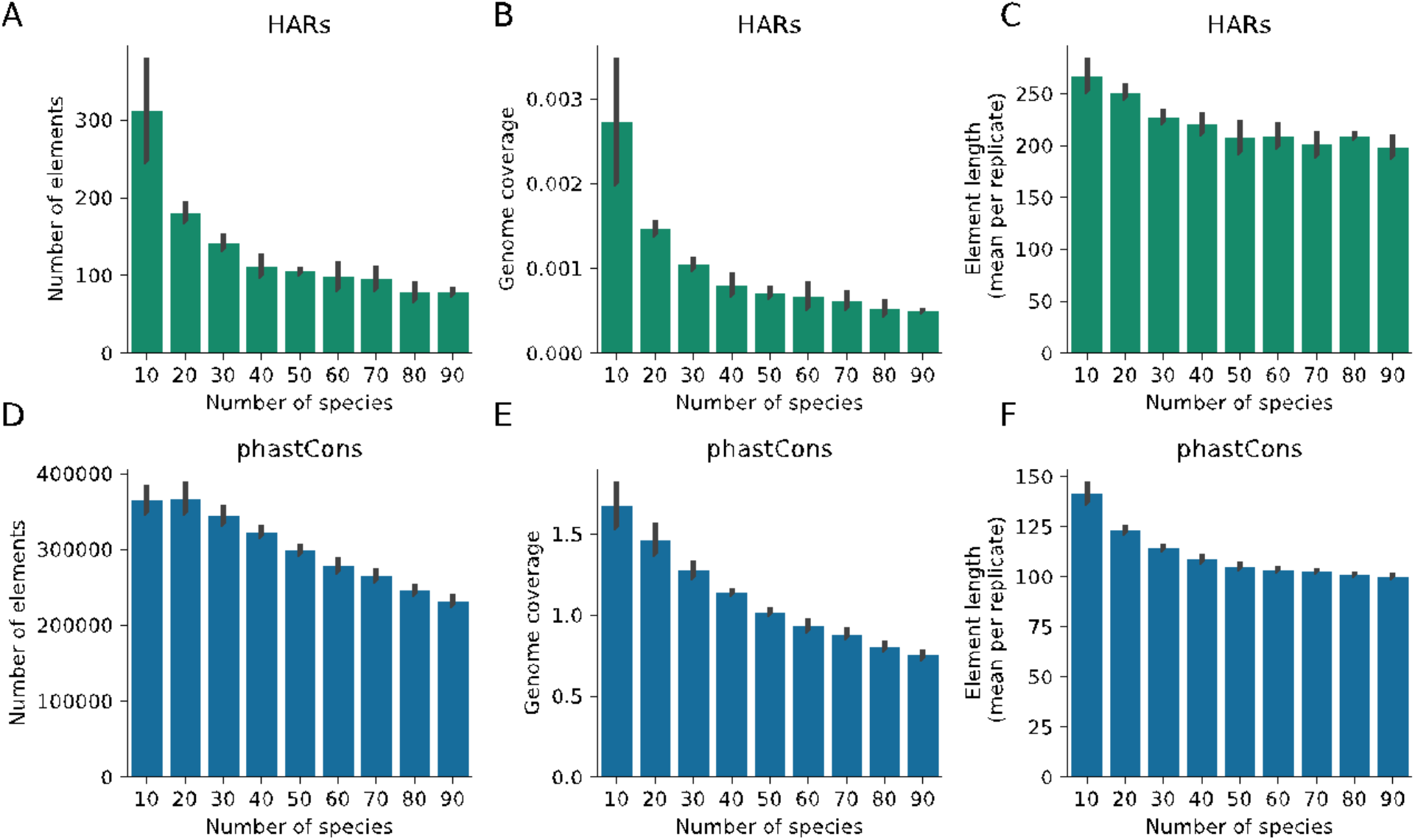
Impact of species number on UCSC HARs and phastCons elements identified. Number of elements, genome coverage, and element length as a function of the number of species included in the analysis for UCSC HARs (A, B, C) and phastCons (D, E, F) based on the UCSC 100-way alignment of vertebrates. Error bars indicate standard deviation based on three sets of random draws of species (see Supplemental Text).

**Fig. S12.**
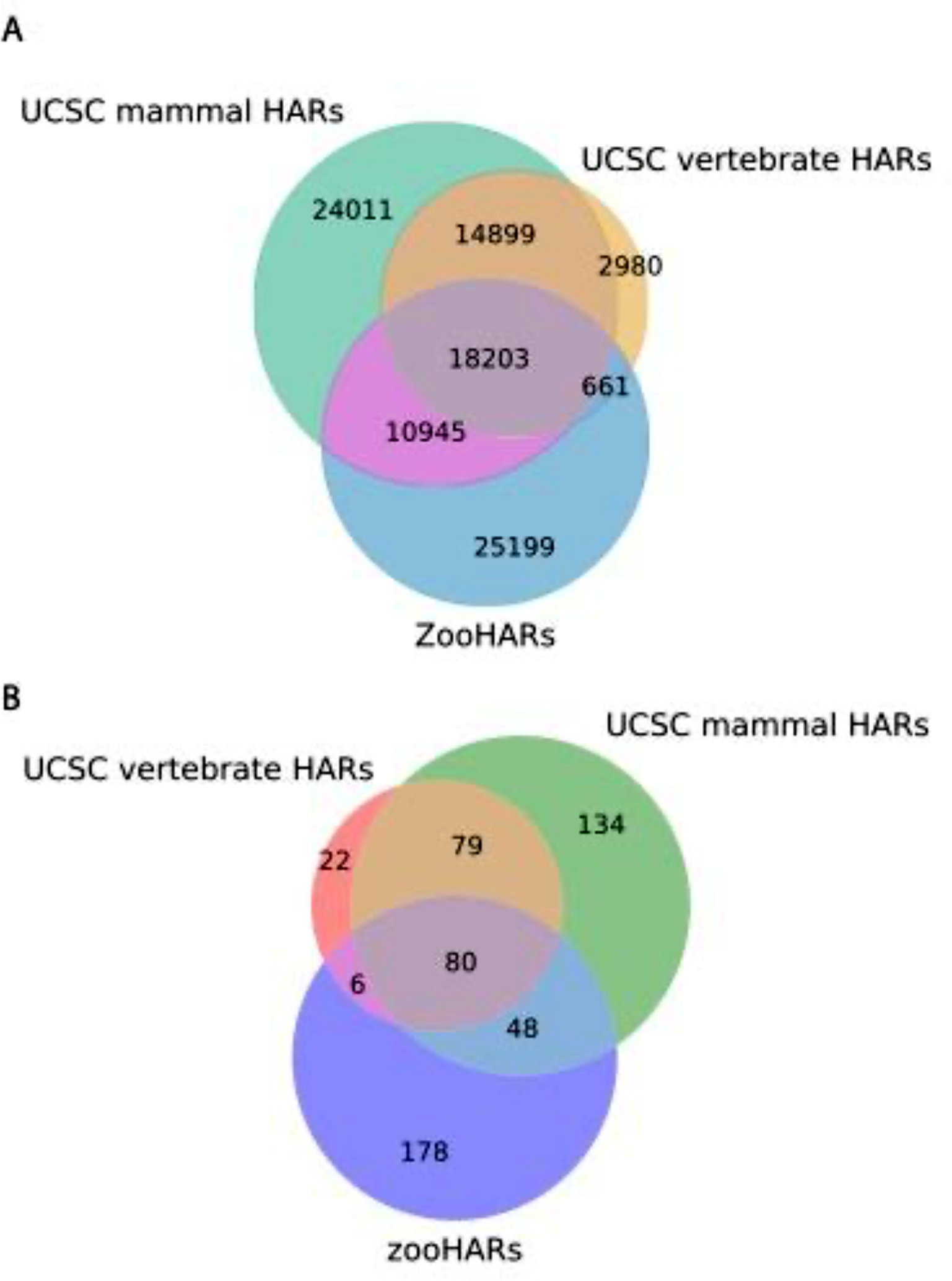
Comparison of Zoonomia with UCSC HARs. Overlap in base pairs (**A**) and elements (**B**) of HARs identified from the entire 100-way UCSC alignment (UCSC vertebrate HARs), the 61-mammal subset of the 100-way UCSC alignment (UCSC mammals HARs), and zooHARs.

**Table S1. (separate file)**

zooHAR coordinates (hg38), selection annotations and enhancer prediction scores.

**Table S2. (separate file)**

zooCHAR coordinates (hg38) and selection annotations.

**Table S3. (separate file)**

Predicted disruption scores for human-specific structural variants.

**Table S4. (separate file)**

Quality control information for the Hi-C data for human and chimpanzee NPCs generated in this work, including sequenced read depths, uniquely mapped pairs and cis/trans ratios for each sample.

**Table S5. (separate file)**

Loops and TADs for human and chimpanzee NPC Hi-C generated in this study.

**Table S6. (separate file)**

zooHAR overlaps with epigenomic annotations and enrichments compared to phastCons elements, and zooHAR activity in an MPRA in primary human mid-gestation telencephalon cells.

**Table S7. (separate file)**

zooHAR and zooCHAR CellWalker assignments based on data from the developing human telencephalon.

## Acknowledgments

We thank Maureen Pittman, Geoffrey Fudenberg, Abigail Lind, Evonne McArthur, Ryan Ziffra, Tony Capra and Svetlana Lyalina for helpful discussions, sharing code and suggestions towards the results shown in this work. We thank Giovanni Maki for assistance with figures and visualization. This project was funded by NIH NHGRI R01HG008742 and the Swedish Research Council Distinguished Professor Award.

## Funding

Discovery Fellowship (KCK)

National Institutes of Health GR-01125 (KCK, KSP)

National Institute of Mental Health R01MH109907, U01MH116438 (NA, KSP)

Gladstone Institutes (KSP)

NIH DP2MH122400-01 (AAP, TF)

Schmidt Futures Foundation (AAP, TF)

Shurl and Kay Curci Foundation (AAP, TF)

NIH NHGRI R01HG008742 (EK)

Swedish Research Council Distinguished Professor Award (KLT)

## Author contributions

Conceptualization: KCK, KSP

Methodology: KCK, SW, PFP, TF, FI, HR, NA, AP, ZC, KSP

Investigation: KCK, SW, PFP, TF, FI, HR Visualization: KCK, PFP

Funding acquisition: NA, KSP

Supervision: NA, AP, KSP

Writing - original draft: KCK, KSP

Writing - review & editing: All authors

## Competing interests

Authors declare they have no competing interests.

## Data and materials availability

The Zoonomia data are available at https://zoonomiaproject.org/the-project/. The Nextflow pipeline to identify lineage-specific accelerated regions is available at https://github.com/keoughkath/AcceleratedRegionsNF. The Hi-C data are available at GSE183137. All other data are available in the main text or the supplementary materials.

## Zoonomia Consortium Authors - Collaborators

Gregory Andrews^1^, Joel C. Armstrong^2^, Matteo Bianchi^3^, Bruce W. Birren^4^, Kevin Bredemeyer^5^, Ana M Breit^6^, Matthew J Christmas^3^, Joana Damas^7^, Mark Diekhans^2^, Michael X. Dong^3^, Eduardo Eizirik^8^, Kaili Fan^1^, Cornelia Fanter^9^, Nicole M. Foley^5^, Karin Forsberg-Nilsson^10^, Carlos J. Garcia^11^, John Gatesy^12^, Steven Gazal^13^, Diane P. Genereux^4^, Daniel Goodman^14^, Linda Goodman^15^, Jenna Grimshaw^11^, Michaela K. Halsey^11^, Andrew Harris^5^, Glenn Hickey^16^, Michael Hiller^17^, Allyson Hindle^9^, Robert M. Hubley^18^, Graham Hughes^19^, Jeremy Johnson^4^, David Juan^20^, Irene M. Kaplow^21,22^, Elinor K. Karlsson^1,4^, Kathleen C. Keough^23,24^, Bogdan Kirilenko^17^, Jennifer M. Korstian^11^, Sergey V. Kozyrev^3^, Alyssa J. Lawler^25^, Colleen Lawless^19^, Danielle L. Levesque^6^, Harris A. Lewin^7,26,27^, Xue Li^1,4^, Abigail Lind^23,24^, Kerstin Lindblad-Toh^3,4^, Voichita D. Marinescu^3^, Tomas Marques-Bonet^20^, Victor Mason^28^, Jennifer R. S. Meadows^3^, Jill Moore^1^, Diana D. Moreno-Santillan^11^, Kathleen M. Morrill^1,4^, Gerhard Muntané^20^, William Murphy^5^, Arcadi Navarro^20^, Martin Nweeia^29,30,31,32^, Austin Osmanski^11^, Benedict Paten^2^, Nicole S. Paulat^11^, Eric Pederson^3^, Andreas R. Pfenning^21,22^, BaDoi N. Phan^21^, Katherine S. Pollard^23,24,33^, Kavya Prasad^21^, Henry Pratt^1^, David A. Ray^11^, Jeb Rosen^18^, Irina Ruf^34^, Louise Ryan^19^, Oliver Ryder^35,36^, Daniel Schäffer^21^, Aitor Serres^20^, Beth Shapiro^37,38^, Arian F. A. Smit^18^, Mark Springer^39^, Chaitanya Srinivasan^21^, Cynthia Steiner^40^, Jessica M. Storer^18^, Patrick F. Sullivan^41^, Kevin A. M. Sullivan^10^, Elisabeth Sundström^3^, Megan A Supple^38^, Ross Swofford^4^, Joy-El Talbot^42^, Emma Teeling^19^, Jason Turner-Maier^4^, Alejandro Valenzuela^20^, Franziska Wagner^34^, Ola Wallerman^3^, Chao Wang^3^, Juehan Wang^13^, Zhiping Weng^1^, Aryn P. Wilder^35^, Morgan E. Wirthlin^21,22^, Shuyang Yao^43^, Xiaomeng Zhang^2^

1 Program in Bioinformatics and Integrative Biology, University of Massachusetts Medical School, Worcester, MA 01605, USA

2 Genomics Institute, UC Santa Cruz, 1156 High Street, Santa Cruz, CA 95064, USA

3 Science for Life Laboratory, Department of Medical Biochemistry and Microbiology, Uppsala University, Uppsala, 751 32, Sweden

4 Broad Institute of MIT and Harvard, Cambridge MA 02139, USA

5 Veterinary Integrative Biosciences, Texas A&M University, College Station, TX 77843, USA

6 School of Biology and Ecology, University of Maine, Orono, Maine 04469, USA

7 The Genome Center, University of California Davis, Davis, CA 95616, USA

8 School of Health and Life Sciences, Pontifical Catholic University of Rio Grande do Sul, Porto Alegre, 90619-900, Brazil

9 School of Life Sciences, University of Nevada Las Vegas, Las Vegas, NV 89154, USA

10 Department of Immunology, Genetics and Pathology, Science for Life Laboratory, Uppsala University, Uppsala, 751 85, Sweden

11 Department of Biological Sciences, Texas Tech University, Lubbock, TX 79409, USA

12 Division of Vertebrate Zoology, American Museum of Natural History, New York, NY 10024, USA

13 Keck School of Medicine, University of Southern California, Los Angeles, CA 90033, USA

14 University of California San Francisco, San Francisco, CA 94143 USA

15 Fauna Bio Inc., Emeryville, CA 94608, USA

16 Baskin School of Engineering, University of California Santa Cruz, Santa Cruz, CA 95064, USA

17 Max Planck Institute of Molecular Cell Biology and Genetics, 01307, Dresden, Germany

18 Institute for Systems Biology, Seattle, WA 98109, USA

19 School of Biology and Environmental Science, University College Dublin, Belfield, Dublin 4, Ireland

20 Institute of Evolutionary Biology (UPF-CSIC), Department of Experimental and Health Sciences, Universitat Pompeu Fabra, Barcelona, 08003, Spain

21 Department of Computational Biology, School of Computer Science, Carnegie Mellon University, Pittsburgh, PA 15213, USA

22 Neuroscience Institute, Carnegie Mellon University, Pittsburgh, PA 15213, USA

23 Gladstone Institutes, San Francisco, CA 94158, USA

24 Department of Epidemiology & Biostatistics, University of California, San Francisco, CA 94158, USA

25 Department of Biology, Carnegie Mellon University, Pittsburgh, PA 15213, USA

26 Department of Evolution and Ecology, University of California, Davis, CA 95616, USA

27 John Muir Institute for the Environment, University of California, Davis, CA 95616, USA

28 Institute of Cell Biology, University of Bern, 3012 Bern, Switzerland

29 Narwhal Genome Initiative, Department of Restorative Dentistry and Biomaterials Sciences, Harvard School of Dental Medicine, Boston, MA 02115, USA

30 Department of Comprehensive Care, School of Dental Medicine, Case Western Reserve University, Cleveland, OH 44106, USA

31 Department of Vertebrate Zoology, Smithsonian Institution, Washington, DC 20002, USA

32 Department of Vertebrate Zoology, Canadian Museum of Nature, Ottawa, Ontario K2P 2R1, Canada

33 Chan Zuckerberg Biohub, San Francisco, CA 94158, USA

34 Division of Messel Research and Mammalogy, Senckenberg Research Institute and Natural History Museum Frankfurt, 60325 Frankfurt am Main, Germany

35 Conservation Genetics, San Diego Zoo Wildlife Alliance, Escondido, CA 92027, USA

36 Department of Evolution, Behavior and Ecology, Division of Biology, University of California, San Diego, La Jolla, CA 92039 USA

37 Howard Hughes Medical Institute, University of California Santa Cruz, Santa Cruz, CA 95064, USA

38 Department of Ecology and Evolutionary Biology, University of California Santa Cruz, Santa Cruz, CA 95064, USA

39 Department of Evolution, Ecology and Organismal Biology, University of California, Riverside, CA 92521, USA

40 Conservation Science Wildlife Health, San Diego Zoo Wildlife Alliance, Escondido CA 92027, USA

41 Department of Genetics, University of North Carolina Medical School, Chapel Hill, NC 27599, USA

42 Iris Data Solutions, LLC, Orono, ME 04473, USA

43 Department of medical Epidemiology and Biostatistics, Karolinska Institute, Stockholm, 171 77, Sweden

## Notes

### Competing Interest Statement

The authors have declared no competing interest.

## References and Notes

1. K. S. Pollard, S. R. Salama, B. King, A. D. Kern, T. Dreszer, S. Katzman, A. Siepel, J. S. Pedersen, G. Bejerano, R. Baertsch, K. R. Rosenbloom, J. Kent, D. Haussler, Forces shaping the fastest evolving regions in the human genome. PLoS Genet. 2, e168 (2006).

2. K. S. Pollard, S. R. Salama, N. Lambert, M.-A. Lambot, S. Coppens, J. S. Pedersen, S. Katzman, B. King, C. Onodera, A. Siepel, A. D. Kern, C. Dehay, H. Igel, M. Ares Jr, P. Vanderhaeghen, D. Haussler, An RNA gene expressed during cortical development evolved rapidly in humans. Nature. 443, 167–172 (2006).

3. S. Prabhakar, J. P. Noonan, S. Pääbo, E. M. Rubin, Accelerated evolution of conserved noncoding sequences in humans. Science. 314, 786 (2006).

4. C. P. Bird, B. E. Stranger, M. Liu, D. J. Thomas, C. E. Ingle, C. Beazley, W. Miller, M. E. Hurles, E. T. Dermitzakis, Fast-evolving noncoding sequences in the human genome. Genome Biol. 8, R118 (2007).

5. E. C. Bush, B. T. Lahn, A genome-wide screen for noncoding elements important in primate evolution. BMC Evol. Biol. 8, 17 (2008).

6. M. J. Hubisz, K. S. Pollard, Exploring the genesis and functions of Human Accelerated Regions sheds light on their role in human evolution. Curr. Opin. Genet. Dev. 29, 15–21 (2014).

7. L. F. Franchini, K. S. Pollard, Human evolution: the non-coding revolution. BMC Biol. 15, 89 (2017).

8. S. K. Reilly, J. P. Noonan, Evolution of Gene Regulation in Humans. Annu. Rev. Genomics Hum. Genet. 17, 45–67 (2016).

9. J. A. Capra, G. D. Erwin, G. McKinsey, J. L. R. Rubenstein, K. S. Pollard, Many human accelerated regions are developmental enhancers. Philos. Trans. R. Soc. Lond. B Biol. Sci. 368, 20130025 (2013).

10. D. Kostka, M. J. Hubisz, A. Siepel, K. S. Pollard, The role of GC-biased gene conversion in shaping the fastest evolving regions of the human genome. Mol. Biol. Evol. 29, 1047–1057 (2012).

11. Chimpanzee Sequencing and Analysis Consortium, Initial sequence of the chimpanzee genome and comparison with the human genome. Nature. 437, 69–87 (2005).

12. D. Hnisz, A. S. Weintraub, D. S. Day, A.-L. Valton, R. O. Bak, C. H. Li, J. Goldmann, B. R. Lajoie, Z. P. Fan, A. A. Sigova, J. Reddy, D. Borges-Rivera, T. I. Lee, R. Jaenisch, M. H. Porteus, J. Dekker, R. A. Young, Activation of proto-oncogenes by disruption of chromosome neighborhoods. Science. 351, 1454–1458 (2016).

13. M. Affer, M. Chesi, W.-D. G. Chen, J. J. Keats, Y. N. Demchenko, A. V. Roschke, S. Van Wier, R. Fonseca, P. L. Bergsagel, W. M. Kuehl, Promiscuous MYC locus rearrangements hijack enhancers but mostly super-enhancers to dysregulate MYC expression in multiple myeloma. Leukemia. 28, 1725–1735 (2014).

14. M. W. Zimmerman, Y. Liu, S. He, A. D. Durbin, B. J. Abraham, J. Easton, Y. Shao, B. Xu, S. Zhu, X. Zhang, Z. Li, N. Weichert-Leahey, R. A. Young, J. Zhang, A. Thomas Look, MYC Drives a Subset of High-Risk Pediatric Neuroblastomas and Is Activated through Mechanisms Including Enhancer Hijacking and Focal Enhancer Amplification. Cancer Discovery. 8 (2018), pp. 320–335.

15. D. G. Lupiáñez, M. Spielmann, S. Mundlos, Breaking TADs: How Alterations of Chromatin Domains Result in Disease. Trends Genet. 32, 225–237 (2016).

16. J. Ibn-Salem, S. Köhler, M. I. Love, H.-R. Chung, N. Huang, M. E. Hurles, M. Haendel, N. L. Washington, D. Smedley, C. J. Mungall, S. E. Lewis, C.-E. Ott, S. Bauer, P. N. Schofield, S. Mundlos, M. Spielmann, P. N. Robinson, Deletions of chromosomal regulatory boundaries are associated with congenital disease. Genome Biol. 15, 423 (2014).

17. M. Franke, D. M. Ibrahim, G. Andrey, W. Schwarzer, V. Heinrich, R. Schöpflin, K. Kraft, R. Kempfer, I. Jerković, W.-L. Chan, M. Spielmann, B. Timmermann, L. Wittler, I. Kurth, P. Cambiaso, O. Zuffardi, G. Houge, L. Lambie, F. Brancati, A. Pombo, M. Vingron, F. Spitz, S. Mundlos, Formation of new chromatin domains determines pathogenicity of genomic duplications. Nature. 538, 265–269 (2016).

18. R. D. Acemel, I. Maeso, J. L. Gómez-Skarmeta, Topologically associated domains: a successful scaffold for the evolution of gene regulation in animals. Wiley Interdisciplinary Reviews: Developmental Biology. 6 (2017), p. e265.

19. I. Maeso, R. D. Acemel, J. L. Gómez-Skarmeta, Cis-regulatory landscapes in development and evolution. Curr. Opin. Genet. Dev. 43, 17–22 (2017).

20. N. Lonfat, D. Duboule, Structure, function and evolution of topologically associating domains (TADs) at HOX loci. FEBS Lett. 589, 2869–2876 (2015).

21. R. D. Acemel, J. J. Tena, I. Irastorza-Azcarate, F. Marlétaz, C. Gómez-Marín, E. de la Calle-Mustienes, S. Bertrand, S. G. Diaz, D. Aldea, J.-M. Aury, S. Mangenot, P. W. H. Holland, D. P. Devos, I. Maeso, H. Escrivá, J. L. Gómez-Skarmeta, A single three-dimensional chromatin compartment in amphioxus indicates a stepwise evolution of vertebrate Hox bimodal regulation. Nat. Genet. 48, 336–341 (2016).

22. Z. N. Kronenberg, I. T. Fiddes, D. Gordon, S. Murali, S. Cantsilieris, O. S. Meyerson, J. G. Underwood, B. J. Nelson, M. J. P. Chaisson, M. L. Dougherty, K. M. Munson, A. R. Hastie, M. Diekhans, F. Hormozdiari, N. Lorusso, K. Hoekzema, R. Qiu, K. Clark, A. Raja, A. E. Welch, M. Sorensen, C. Baker, R. S. Fulton, J. Armstrong, T. A. Graves-Lindsay, A. M. Denli, E. R. Hoppe, P. Hsieh, C. M. Hill, A. W. C. Pang, J. Lee, E. T. Lam, S. K. Dutcher, F. H. Gage, W. C. Warren, J. Shendure, D. Haussler, V. A. Schneider, H. Cao, M. Ventura, R. K. Wilson, B. Paten, A. Pollen, E. E. Eichler, High-resolution comparative analysis of great ape genomes. Science. 360 (2018), doi:10.1126/science.aar6343.

23. Genereux, Zoonomia consortium., Evolutionary innovation of eutherian mammals. In preparation/submission (Flagship 1).

24. M. C. Marchetto, B. Hrvoj-Mihic, B. E. Kerman, D. X. Yu, K. C. Vadodaria, S. B. Linker, I. Narvaiza, R. Santos, A. M. Denli, A. P. Mendes, R. Oefner, J. Cook, L. McHenry, J. M. Grasmick, K. Heard, C. Fredlender, L. Randolph-Moore, R. Kshirsagar, R. Xenitopoulos, G. Chou, N. Hah, A. R. Muotri, K. Padmanabhan, K. Semendeferi, F. H. Gage, Species-specific maturation profiles of human, chimpanzee and bonobo neural cells. Elife. 8 (2019), doi:10.7554/eLife.37527.

25. A. A. Pollen, A. Bhaduri, M. G. Andrews, T. J. Nowakowski, O. S. Meyerson, M. A. Mostajo-Radji, E. Di Lullo, B. Alvarado, M. Bedolli, M. L. Dougherty, I. T. Fiddes, Z. N. Kronenberg, J. Shuga, A. A. Leyrat, J. A. West, M. Bershteyn, C. B. Lowe, B. J. Pavlovic, S. R. Salama, D. Haussler, E. E. Eichler, A. R. Kriegstein, Establishing Cerebral Organoids as Models of Human-Specific Brain Evolution. Cell. 176, 743–756.e17 (2019).

26. P. Di Tommaso, M. Chatzou, E. W. Floden, P. P. Barja, E. Palumbo, C. Notredame, Nextflow enables reproducible computational workflows. Nat. Biotechnol. 35, 316–319 (2017).

27. M. J. Hubisz, K. S. Pollard, A. Siepel, PHAST and RPHAST: phylogenetic analysis with space/time models. Brief. Bioinform. 12, 41–51 (2011).

28. A. Siepel, K. S. Pollard, D. Haussler, New Methods for Detecting Lineage-Specific Selection. Lecture Notes in Computer Science (2006), pp. 190–205.

29. K. S. Pollard, M. J. Hubisz, K. R. Rosenbloom, A. Siepel, Detection of nonneutral substitution rates on mammalian phylogenies. Genome Res. 20, 110–121 (2010).

30. D. Kostka, A. K. Holloway, K. S. Pollard, Developmental Loci Harbor Clusters of Accelerated Regions That Evolved Independently in Ape Lineages. Mol. Biol. Evol. 35, 2034–2045 (2018).

31. C. Y. McLean, D. Bristor, M. Hiller, S. L. Clarke, B. T. Schaar, C. B. Lowe, A. M. Wenger, G. Bejerano, GREAT improves functional interpretation of cis-regulatory regions. Nat. Biotechnol. 28, 495–501 (2010).

32. C. S. Greene, A. Krishnan, A. K. Wong, E. Ricciotti, R. A. Zelaya, D. S. Himmelstein, R. Zhang, B. M. Hartmann, E. Zaslavsky, S. C. Sealfon, D. I. Chasman, G. A. FitzGerald, K. Dolinski, T. Grosser, O. G. Troyanskaya, Understanding multicellular function and disease with human tissue-specific networks. Nat. Genet. 47, 569–576 (2015).

33. D. Kostka, M. W. Hahn, K. S. Pollard, Noncoding sequences near duplicated genes evolve rapidly. Genome Biol. Evol. 2, 518–533 (2010).

34. S. S. P. Rao, M. H. Huntley, N. C. Durand, E. K. Stamenova, I. D. Bochkov, J. T. Robinson, A. L. Sanborn, I. Machol, A. D. Omer, E. S. Lander, E. L. Aiden, A 3D map of the human genome at kilobase resolution reveals principles of chromatin looping. Cell. 159, 1665–1680 (2014).

35. M. Spielmann, D. G. Lupiáñez, S. Mundlos, Structural variation in the 3D genome. Nat. Rev. Genet. 19, 453–467 (2018).

36. G. Fudenberg, D. R. Kelley, K. S. Pollard, Predicting 3D genome folding from DNA sequence with Akita. Nat. Methods. 17, 1111–1117 (2020).

37. S. Whalen, F. Inoue, H. Ryu, T. Fair, E. Markenscoff-Papadimitriou, K. Keough, M. Kircher, B. Martin, B. Alvarado, O. Elor, D. L. Cintron, A. Williams, M. A. H. Samee, S. Thomas, R. Krencik, E. M. Ullian, A. R. Kriegstein, J. Shendure, A. Pollen, N. Ahituv, K. S. Pollard, Machine-Learning Dissection of Human Accelerated Regions in Primate Neurodevelopment (2022), (available at https://papers.ssrn.com/abstract=4149954).

38. T. Yang, F. Zhang, G. G. Yardımcı, F. Song, R. C. Hardison, W. S. Noble, F. Yue, Q. Li, HiCRep: assessing the reproducibility of Hi-C data using a stratum-adjusted correlation coefficient. Genome Res. 27, 1939–1949 (2017).

39. J. R. Dixon, S. Selvaraj, F. Yue, A. Kim, Y. Li, Y. Shen, M. Hu, J. S. Liu, B. Ren, Topological domains in mammalian genomes identified by analysis of chromatin interactions. Nature. 485, 376–380 (2012).

40. M. Vietri Rudan, M. V. Rudan, C. Barrington, S. Henderson, C. Ernst, D. T. Odom, A. Tanay, S. Hadjur, Comparative Hi-C Reveals that CTCF Underlies Evolution of Chromosomal Domain Architecture. Cell Reports. 10 (2015), pp. 1297–1309.

41. I. E. Eres, K. Luo, C. J. Hsiao, L. E. Blake, Y. Gilad, Reorganization of 3D genome structure may contribute to gene regulatory evolution in primates. PLoS Genet. 15, e1008278 (2019).

42. X. Luo, Y. Liu, D. Dang, T. Hu, Y. Hou, X. Meng, F. Zhang, T. Li, C. Wang, M. Li, H. Wu, Q. Shen, Y. Hu, X. Zeng, X. He, L. Yan, S. Zhang, C. Li, B. Su, 3D Genome of macaque fetal brain reveals evolutionary innovations during primate corticogenesis. Cell. 184, 723–740.e21 (2021).

43. I. E. Eres, Y. Gilad, A TAD Skeptic: Is 3D Genome Topology Conserved? Trends Genet. 37, 216–223 (2021).

44. C. Hoencamp, O. Dudchenko, A. M. O. Elbatsh, S. Brahmachari, J. A. Raaijmakers, T. van Schaik, Á. Sedeño Cacciatore, V. G. Contessoto, R. G. H. P. van Heesbeen, B. van den Broek, A. N. Mhaskar, H. Teunissen, B. G. St Hilaire, D. Weisz, A. D. Omer, M. Pham, Z. Colaric, Z. Yang, S. S. P. Rao, N. Mitra, C. Lui, W. Yao, R. Khan, L. L. Moroz, A. Kohn, J. St Leger, A. Mena, K. Holcroft, M. C. Gambetta, F. Lim, E. Farley, N. Stein, A. Haddad, D. Chauss, A. S. Mutlu, M. C. Wang, N. D. Young, E. Hildebrandt, H. H. Cheng, C. J. Knight, T. L. U. Burnham, K. A. Hovel, A. J. Beel, P.-J. Mattei, R. D. Kornberg, W. C. Warren, G. Cary, J. L. Gómez-Skarmeta, V. Hinman, K. Lindblad-Toh, F. Di Palma, K. Maeshima, A. S. Multani, S. Pathak, L. Nel-Themaat, R. R. Behringer, P. Kaur, R. H. Medema, B. van Steensel, E. de Wit, J. N. Onuchic, M. Di Pierro, E. Lieberman Aiden, B. D. Rowland, 3D genomics across the tree of life reveals condensin II as a determinant of architecture type. Science. 372, 984–989 (2021).

45. L. A. Mirny, M. Imakaev, N. Abdennur, Two major mechanisms of chromosome organization. Current Opinion in Cell Biology. 58 (2019), pp. 142–152.

46. A. Roayaei Ardakany, H. T. Gezer, S. Lonardi, F. Ay, Mustache: multi-scale detection of chromatin loops from Hi-C and Micro-C maps using scale-space representation. Genome Biol. 21, 256 (2020).

47. A. G. Diehl, N. Ouyang, A. P. Boyle, Transposable elements contribute to cell and species-specific chromatin looping and gene regulation in mammalian genomes. Nat. Commun. 11, 1796 (2020).

48. S. Kanton, M. J. Boyle, Z. He, M. Santel, A. Weigert, F. Sanchís-Calleja, P. Guijarro, L. Sidow, J. S. Fleck, D. Han, Z. Qian, M. Heide, W. B. Huttner, P. Khaitovich, S. Pääbo, B. Treutlein, J. G. Camp, Organoid single-cell genomic atlas uncovers human-specific features of brain development. Nature. 574, 418–422 (2019).

49. B. J. Pavlovic, L. E. Blake, J. Roux, C. Chavarria, Y. Gilad, A Comparative Assessment of Human and Chimpanzee iPSC-derived Cardiomyocytes with Primary Heart Tissues. Sci. Rep. 8, 15312 (2018).

50. S. K. Sundaram, A. M. Huq, Z. Sun, W. Yu, L. Bennett, B. J. Wilson, M. E. Behen, H. T. Chugani, Exome sequencing of a pedigree with tourette syndrome or chronic tic disorder. Annals of Neurology. 69 (2011), pp. 901–904.

51. K. Lindblad-Toh, M. Garber, O. Zuk, M. F. Lin, B. J. Parker, S. Washietl, P. Kheradpour, J. Ernst, G. Jordan, E. Mauceli, L. D. Ward, C. B. Lowe, A. K. Holloway, M. Clamp, S. Gnerre, J. Alföldi, K. Beal, J. Chang, H. Clawson, J. Cuff, F. Di Palma, S. Fitzgerald, P. Flicek, M. Guttman, M. J. Hubisz, D. B. Jaffe, I. Jungreis, W. J. Kent, D. Kostka, M. Lara, A. L. Martins, T. Massingham, I. Moltke, B. J. Raney, M. D. Rasmussen, J. Robinson, A. Stark, A. J. Vilella, J. Wen, X. Xie, M. C. Zody, Broad Institute Sequencing Platform and Whole Genome Assembly Team, J. Baldwin, T. Bloom, C. W. Chin, D. Heiman, R. Nicol, C. Nusbaum, S. Young, J. Wilkinson, K. C. Worley, C. L. Kovar, D. M. Muzny, R. A. Gibbs, Baylor College of Medicine Human Genome Sequencing Center Sequencing Team, A. Cree, H. H. Dihn, G. Fowler, S. Jhangiani, V. Joshi, S. Lee, L. R. Lewis, L. V. Nazareth, G. Okwuonu, J. Santibanez, W. C. Warren, E. R. Mardis, G. M. Weinstock, R. K. Wilson, Genome Institute at Washington University, K. Delehaunty, D. Dooling, C. Fronik, L. Fulton, B. Fulton, T. Graves, P. Minx, E. Sodergren, E. Birney, E. H. Margulies, J. Herrero, E. D. Green, D. Haussler, A. Siepel, N. Goldman, K. S. Pollard, J. S. Pedersen, E. S. Lander, M. Kellis, A high-resolution map of human evolutionary constraint using 29 mammals. Nature. 478, 476–482 (2011).

52. R. N. Doan, B.-I. Bae, B. Cubelos, C. Chang, A. A. Hossain, S. Al-Saad, N. M. Mukaddes, O. Oner, M. Al-Saffar, S. Balkhy, G. G. Gascon, Homozygosity Mapping Consortium for Autism, M. Nieto, C. A. Walsh, Mutations in Human Accelerated Regions Disrupt Cognition and Social Behavior. Cell. 167, 341–354.e12 (2016).

53. B. Castelijns, M. L. Baak, I. S. Timpanaro, C. R. M. Wiggers, M. W. Vermunt, P. Shang, I. Kondova, G. Geeven, V. Bianchi, W. de Laat, N. Geijsen, M. P. Creyghton, Hominin-specific regulatory elements selectively emerged in oligodendrocytes and are disrupted in autism patients. Nat. Commun. 11, 301 (2020).

54. E. Markenscoff-Papadimitriou, S. Whalen, P. Przytycki, R. Thomas, F. Binyameen, T. J. Nowakowski, A. R. Kriegstein, S. J. Sanders, M. W. State, K. S. Pollard, J. L. Rubenstein, A Chromatin Accessibility Atlas of the Developing Human Telencephalon. Cell. 182, 754–769.e18 (2020).

55. M. P. Forrest, H. Zhang, W. Moy, H. McGowan, C. Leites, L. E. Dionisio, Z. Xu, J. Shi, A. R. Sanders, W. J. Greenleaf, C. A. Cowan, Z. P. Pang, P. V. Gejman, P. Penzes, J. Duan, Open Chromatin Profiling in hiPSC-Derived Neurons Prioritizes Functional Noncoding Psychiatric Risk Variants and Highlights Neurodevelopmental Loci. Cell Stem Cell. 21, 305–318.e8 (2017).

56. C. C. Funk, A. M. Casella, S. Jung, M. A. Richards, A. Rodriguez, P. Shannon, R. Donovan-Maiye, B. Heavner, K. Chard, Y. Xiao, G. Glusman, N. Ertekin-Taner, T. E. Golde, A. Toga, L. Hood, J. D. Van Horn, C. Kesselman, I. Foster, R. Madduri, N. D. Price, S. A. Ament, Atlas of Transcription Factor Binding Sites from ENCODE DNase Hypersensitivity Data across 27 Tissue Types. Cell Rep. 32, 108029 (2020).

57. M. Song, X. Yang, X. Ren, L. Maliskova, B. Li, I. R. Jones, C. Wang, F. Jacob, K. Wu, M. Traglia, T. W. Tam, K. Jamieson, S.-Y. Lu, G.-L. Ming, Y. Li, J. Yao, L. A. Weiss, J. R. Dixon, L. M. Judge, B. R. Conklin, H. Song, L. Gan, Y. Shen, Mapping cis-regulatory chromatin contacts in neural cells links neuropsychiatric disorder risk variants to target genes. Nat. Genet. 51, 1252–1262 (2019).

58. M. Song, M.-P. Pebworth, X. Yang, A. Abnousi, C. Fan, J. Wen, J. D. Rosen, M. N. K. Choudhary, X. Cui, I. R. Jones, S. Bergenholtz, U. C. Eze, I. Juric, B. Li, L. Maliskova, J. Lee, W. Liu, A. A. Pollen, Y. Li, T. Wang, M. Hu, A. R. Kriegstein, Y. Shen, Cell-type-specific 3D epigenomes in the developing human cortex. Nature. 587, 644–649 (2020).

59. R. S. Ziffra, C. N. Kim, J. M. Ross, A. Wilfert, T. N. Turner, M. Haeussler, A. M. Casella, P. F. Przytycki, K. C. Keough, D. Shin, D. Bogdanoff, A. Kreimer, K. S. Pollard, S. A. Ament, E. E. Eichler, N. Ahituv, T. J. Nowakowski, Single-cell epigenomics reveals mechanisms of human cortical development. Nature. 598, 205–213 (2021).

60. Roadmap Epigenomics Consortium, A. Kundaje, W. Meuleman, J. Ernst, M. Bilenky, A. Yen, A. Heravi-Moussavi, P. Kheradpour, Z. Zhang, J. Wang, M. J. Ziller, V. Amin, J. W. Whitaker, M. D. Schultz, L. D. Ward, A. Sarkar, G. Quon, R. S. Sandstrom, M. L. Eaton, Y.-C. Wu, A. R. Pfenning, X. Wang, M. Claussnitzer, Y. Liu, C. Coarfa, R. A. Harris, N. Shoresh, C. B. Epstein, E. Gjoneska, D. Leung, W. Xie, R. D. Hawkins, R. Lister, C. Hong, P. Gascard, A. J. Mungall, R. Moore, E. Chuah, A. Tam, T. K. Canfield, R. S. Hansen, R. Kaul, P. J. Sabo, M. S. Bansal, A. Carles, J. R. Dixon, K.-H. Farh, S. Feizi, R. Karlic, A.-R. Kim, A. Kulkarni, D. Li, R. Lowdon, G. Elliott, T. R. Mercer, S. J. Neph, V. Onuchic, P. Polak, N. Rajagopal, P. Ray, R. C. Sallari, K. T. Siebenthall, N. A. Sinnott-Armstrong, M. Stevens, R. E. Thurman, J. Wu, B. Zhang, X. Zhou, A. E. Beaudet, L. A. Boyer, P. L. De Jager, P. J. Farnham, S. J. Fisher, D. Haussler, S. J. M. Jones, W. Li, M. A. Marra, M. T. McManus, S. Sunyaev, J. A. Thomson, T. D. Tlsty, L.-H. Tsai, W. Wang, R. A. Waterland, M. Q. Zhang, L. H. Chadwick, B. E. Bernstein, J. F. Costello, J. R. Ecker, M. Hirst, A. Meissner, A. Milosavljevic, B. Ren, J. A. Stamatoyannopoulos, T. Wang, M. Kellis, Integrative analysis of 111 reference human epigenomes. Nature. 518, 317–330 (2015).

61. M. A. Petryniak, G. B. Potter, D. H. Rowitch, J. L. R. Rubenstein, Dlx1 and Dlx2 control neuronal versus oligodendroglial cell fate acquisition in the developing forebrain. Neuron. 55, 417–433 (2007).

62. M. T. Schmitz, K. Sandoval, C. P. Chen, M. A. Mostajo-Radji, W. W. Seeley, T. J. Nowakowski, C. J. Ye, M. F. Paredes, A. A. Pollen, The development and evolution of inhibitory neurons in primate cerebrum. Nature. 603, 871–877 (2022).

63. M. Shibata, K. Pattabiraman, B. Lorente-Galdos, D. Andrijevic, X. Xing, A. M. M. Sousa, G. Santpere, N. Sestan, Regulation of Prefrontal Patterning, Connectivity and Synaptogenesis by Retinoic Acid. bioRxiv (2019), p. 2019.12.31.891036.

64. A. Visel, S. Minovitsky, I. Dubchak, L. A. Pennacchio, VISTA Enhancer Browser—a database of tissue-specific human enhancers. Nucleic Acids Res. 35, D88–D92 (2006).

65. P. F. Przytycki, K. S. Pollard, CellWalker integrates single-cell and bulk data to resolve regulatory elements across cell types in complex tissues. Genome Biol. 22, 61 (2021).

66. P. F. Przytycki, K. S. Pollard, CellWalkR: An R Package for integrating and visualizing single-cell and bulk data to resolve regulatory elements. Bioinformatics (2022), doi:10.1093/bioinformatics/btac150.

67. T. J. Nowakowski, A. Bhaduri, A. A. Pollen, B. Alvarado, M. A. Mostajo-Radji, E. Di Lullo, M. Haeussler, C. Sandoval-Espinosa, S. J. Liu, D. Velmeshev, J. R. Ounadjela, J. Shuga, X. Wang, D. A. Lim, J. A. West, A. A. Leyrat, W. J. Kent, A. R. Kriegstein, Spatiotemporal gene expression trajectories reveal developmental hierarchies of the human cortex. Science. 358, 1318–1323 (2017).

68. R. D. Hodge, T. E. Bakken, J. A. Miller, K. A. Smith, E. R. Barkan, L. T. Graybuck, J. L. Close, B. Long, N. Johansen, O. Penn, Z. Yao, J. Eggermont, T. Höllt, B. P. Levi, S. I. Shehata, B. Aevermann, A. Beller, D. Bertagnolli, K. Brouner, T. Casper, C. Cobbs, R. Dalley, N. Dee, S.-L. Ding, R. G. Ellenbogen, O. Fong, E. Garren, J. Goldy, R. P. Gwinn, D. Hirschstein, C. D. Keene, M. Keshk, A. L. Ko, K. Lathia, A. Mahfouz, Z. Maltzer, M. McGraw, T. N. Nguyen, J. Nyhus, J. G. Ojemann, A. Oldre, S. Parry, S. Reynolds, C. Rimorin, N. V. Shapovalova, S. Somasundaram, A. Szafer, E. R. Thomsen, M. Tieu, G. Quon, R. H. Scheuermann, R. Yuste, S. M. Sunkin, B. Lelieveldt, D. Feng, L. Ng, A. Bernard, M. Hawrylycz, J. W. Phillips, B. Tasic, H. Zeng, A. R. Jones, C. Koch, E. S. Lein, Conserved cell types with divergent features in human versus mouse cortex. Nature. 573, 61–68 (2019).

69. B. Tasic, Z. Yao, L. T. Graybuck, K. A. Smith, T. N. Nguyen, D. Bertagnolli, J. Goldy, E. Garren, M. N. Economo, S. Viswanathan, O. Penn, T. Bakken, V. Menon, J. Miller, O. Fong, K. E. Hirokawa, K. Lathia, C. Rimorin, M. Tieu, R. Larsen, T. Casper, E. Barkan, M. Kroll, S. Parry, N. V. Shapovalova, D. Hirschstein, J. Pendergraft, H. A. Sullivan, T. K. Kim, A. Szafer, N. Dee, P. Groblewski, I. Wickersham, A. Cetin, J. A. Harris, B. P. Levi, S. M. Sunkin, L. Madisen, T. L. Daigle, L. Looger, A. Bernard, J. Phillips, E. Lein, M. Hawrylycz, K. Svoboda, A. R. Jones, C. Koch, H. Zeng, Shared and distinct transcriptomic cell types across neocortical areas. Nature. 563, 72–78 (2018).

70. J. D. Hocker, O. B. Poirion, F. Zhu, J. Buchanan, K. Zhang, J. Chiou, T.-M. Wang, Q. Zhang, X. Hou, Y. E. Li, Y. Zhang, E. N. Farah, A. Wang, A. D. McCulloch, K. J. Gaulton, B. Ren, N. C. Chi, S. Preissl, Cardiac cell type–specific gene regulatory programs and disease risk association. Science Advances. 7, eabf1444 (2021).

71. H. Won, J. Huang, C. K. Opland, C. L. Hartl, D. H. Geschwind, Human evolved regulatory elements modulate genes involved in cortical expansion and neurodevelopmental disease susceptibility. Nat. Commun. 10, 2396 (2019).

72. A. Siepel, G. Bejerano, J. S. Pedersen, A. S. Hinrichs, M. Hou, K. Rosenbloom, H. Clawson, J. Spieth, L. W. Hillier, S. Richards, G. M. Weinstock, R. K. Wilson, R. A. Gibbs, W. J. Kent, W. Miller, D. Haussler, Evolutionarily conserved elements in vertebrate, insect, worm, and yeast genomes. Genome Res. 15, 1034–1050 (2005).

73. D. Karolchik, A. S. Hinrichs, T. S. Furey, K. M. Roskin, C. W. Sugnet, D. Haussler, W. J. Kent, The UCSC Table Browser data retrieval tool. Nucleic Acids Res. 32, D493–6 (2004).

74. A. Frankish, M. Diekhans, A.-M. Ferreira, R. Johnson, I. Jungreis, J. Loveland, J. M. Mudge, C. Sisu, J. Wright, J. Armstrong, I. Barnes, A. Berry, A. Bignell, S. C. Sala, J. Chrast, F. Cunningham, T. Di Domenico, S. Donaldson, I. T. Fiddes, C. G. Girón, J. M. Gonzalez, T. Grego, M. Hardy, T. Hourlier, T. Hunt, O. G. Izuogu, J. Lagarde, F. J. Martin, L. Martínez, S. Mohanan, P. Muir, F. C. P. Navarro, A. Parker, B. Pei, F. Pozo, M. Ruffier, B. M. Schmitt, E. Stapleton, M.-M. Suner, I. Sycheva, B. Uszczynska-Ratajczak, J. Xu, A. Yates, D. Zerbino, Y. Zhang, B. Aken, J. S. Choudhary, M. Gerstein, R. Guigó, T. J. P. Hubbard, M. Kellis, B. Paten, A. Reymond, M. L. Tress, P. Flicek, GENCODE reference annotation for the human and mouse genomes. Nucleic Acids Research. 47 (2019), pp. D766–D773.

75. X. Jiao, B. T. Sherman, D. W. Huang, R. Stephens, M. W. Baseler, H. C. Lane, R. A. Lempicki, DAVID-WS: a stateful web service to facilitate gene/protein list analysis. Bioinformatics. 28, 1805–1806 (2012).

76. S. J. Liu, T. J. Nowakowski, A. A. Pollen, J. H. Lui, M. A. Horlbeck, F. J. Attenello, D. He, J. S. Weissman, A. R. Kriegstein, A. A. Diaz, D. A. Lim, Single-cell analysis of long non-coding RNAs in the developing human neocortex. Genome Biol. 17, 67 (2016).

77. A. S. Hinrichs, D. Karolchik, R. Baertsch, G. P. Barber, G. Bejerano, H. Clawson, M. Diekhans, T. S. Furey, R. A. Harte, F. Hsu, J. Hillman-Jackson, R. M. Kuhn, J. S. Pedersen, A. Pohl, B. J. Raney, K. R. Rosenbloom, A. Siepel, K. E. Smith, C. W. Sugnet, A. Sultan-Qurraie, D. J. Thomas, H. Trumbower, R. J. Weber, M. Weirauch, A. S. Zweig, D. Haussler, W. J. Kent, The UCSC Genome Browser Database: update 2006. Nucleic Acids Res. 34, D590–8 (2006).

78. O. Tange, GNU Parallel 20200922 (‘Ginsburg’) (2020), doi:10.5281/zenodo.4045386.

79. S. Venev, N. Abdennur, A. Goloborodko, I. Flyamer, G. Fudenberg, J. Nuebler, A. Galitsyna, B. Akgol, S. Abraham, P. Kerpedjiev, M. Imakaev, mirnylab/cooltools: v0.3.2 (Zenodo, 2020; https://zenodo.org/record/3787004).

80. H. Li, Aligning sequence reads, clone sequences and assembly contigs with BWA-MEM (2013), (available at http://arxiv.org/abs/1303.3997).

81. open2c, open2c/pairtools, (available at https://github.com/open2c/pairtools).

82. M. Imakaev, G. Fudenberg, R. P. McCord, N. Naumova, A. Goloborodko, B. R. Lajoie, J. Dekker, L. A. Mirny, Iterative correction of Hi-C data reveals hallmarks of chromosome organization. Nat. Methods. 9, 999–1003 (2012).

83. B. R. Lajoie, J. Dekker, N. Kaplan, The Hitchhiker’s guide to Hi-C analysis: Practical guidelines. Methods. 72 (2015), pp. 65–75.

84. open2c, open2c/open2c_examples, (available at https://github.com/open2c/open2c_examples).

85. M. G. Gordon, F. Inoue, B. Martin, M. Schubach, V. Agarwal, S. Whalen, S. Feng, J. Zhao, T. Ashuach, R. Ziffra, A. Kreimer, I. Georgakopoulos-Soares, N. Yosef, C. J. Ye, K. S. Pollard, J. Shendure, M. Kircher, N. Ahituv, lentiMPRA and MPRAflow for high-throughput functional characterization of gene regulatory elements. Nat. Protoc. 15, 2387–2412 (2020).

86. L. Duret, N. Galtier, Biased gene conversion and the evolution of mammalian genomic landscapes. Annu. Rev. Genomics Hum. Genet. 10, 285–311 (2009).

87. E. Takao-Rikitsu, S. Mochida, E. Inoue, M. Deguchi-Tawarada, M. Inoue, T. Ohtsuka, Y. Takai, Physical and functional interaction of the active zone proteins, CAST, RIM1, and Bassoon, in neurotransmitter release. J. Cell Biol. 164, 301–311 (2004).

88. M. P. Coba, N. H. Komiyama, J. Nithianantharajah, M. V. Kopanitsa, T. Indersmitten, N. G. Skene, E. J. Tuck, D. G. Fricker, K. A. Elsegood, L. E. Stanford, N. O. Afinowi, L. M. Saksida, T. J. Bussey, T. J. O’Dell, S. G. N. Grant, TNiK is required for postsynaptic and nuclear signaling pathways and cognitive function. J. Neurosci. 32, 13987–13999 (2012).

89. H. Won, L. de la Torre-Ubieta, J. L. Stein, N. N. Parikshak, J. Huang, C. K. Opland, M. J. Gandal, G. J. Sutton, F. Hormozdiari, D. Lu, C. Lee, E. Eskin, I. Voineagu, J. Ernst, D. H. Geschwind, Chromosome conformation elucidates regulatory relationships in developing human brain. Nature. 538, 523–527 (2016).

